# Self-cyclisation as a general and efficient platform for peptide and protein macrocyclisation

**DOI:** 10.1101/2022.07.12.499226

**Authors:** Xinying Jia, Yanni K.-Y. Chin, Alan Zhang, Theo Crawford, Yifei Zhu, Nicholas L. Fletcher, Zihan Zhou, Brett R. Hamilton, Martin Stroet, Kristofer J. Thurecht, Mehdi Mobli

## Abstract

Macrocyclisation of proteins and peptides results in a remarkable increase in structural stability, making cyclic peptides and proteins of great interest in drug discovery—either directly as drug leads or as in the case of cyclised nanodiscs (cNDs), as tools for studies of trans-membrane receptors and membrane-active peptides. Various biological methods have been developed that are capable of yielding head-to-tail macrocyclised products. Such enzymatic methods require careful optimisation of cyclisation over polymerisation. Here, we describe the engineering of self-cyclising “*autocyclase*” proteins, where an intramolecular rearrangement can be triggered to yield a monomeric cyclic product in high yields. We characterise the self-cyclisation reaction mechanism and demonstrate how the unimolecular reaction path can circumvent existing challenges of enzymatic cyclisation. We use the method to produce several notable cyclic peptides and proteins, demonstrating how autocyclases offer a simple and scalable way to access a vast diversity of macrocyclic biomolecules.

## Introduction

Head-to-tail macrocyclisation is a naturally occurring post-translational modification that stabilises the protein fold, leading to enhanced thermal stability and resistance to proteolytic digestion by exoproteases^1-4^. Mimicry of this natural phenomenon has and continues to inspire protein engineering efforts^5, 6^ including macrocyclic peptide drug leads^7^ and highly stable cyclised lipid nanodiscs used in biophysical characterisation of membrane proteins and membrane active peptides^8, 9^.

The most common strategy to achieve macrocyclisation is by ligation of the termini of the peptide chain through a peptide bond. Chemically this can be achieved efficiently through native chemical ligation (NCL)^10^. NCL, however, becomes challenging for peptides and proteins longer than 100 amino acids. Consequently, several size-insensitive biological approaches have been developed, such as expressed protein ligation (EPL)^11-14^, split-intein mediated protein trans-splicing^15^, and genetic-code reprogramming^16^. EPL and split-intein mediated protein cyclisation both require a free cysteine in the sequence to perform an N-S acyl shift and trans-thioesterification, while backbone cyclisation via codon reprogramming requires the introduction of at least one nonproteinogenic amino acid.

The backbones of peptides and proteins are naturally cyclised by a certain group of proteases with unusual enzymatic transpeptidation activity (as a cyclase or ligase). To date, four stand-alone and ATP-independent ligases have been identified: (*i*) the bacterial transpeptidases, including sortase A^17, 18^; (*ii*) ligases derived from trypsin (trypsiligase)^19^; (*iii*) ligases derived from subtilisin (subtiligase^20^, peptiligase^21^, omniligase-1^22^); and (*iv*) a number of plant-derived ligase-type asparaginyl endopeptidases (AEPs)^23-26^ including butelase-1^27^ and a number of OaAEPs^28 29, 30^. Of these, the bacterial enzyme, sortase A (SrtA)^17, 18^ is perhaps the most popular ligase for peptide and protein cyclisation.

Enzymatic protein or peptide cyclisation reactions are achieved by a conventional bimolecular reaction involving the polypeptide substrate and the enzyme, and while very powerful, these bimolecular reactions involve competition between the cyclisation reaction and a polymerisation reaction (joining the tail of one protein to the head of another)^31, 32^. Minimising the risk of polymerisation can be achieved by lowering the protein concentration, but doing so also results in a quadratic drop in cyclisation rates. This makes the overall process challenging to scale up^9, 31^. Indeed, optimisation of protein and enzyme concentrations remains an important and time-consuming step in enzymatic ligation reactions—leading to various innovations in reaction/reactor design^8, 31^. These innovations, however, do not address the root of the polymerisation problem, which is the dual constraint of maximising a desired diffusion-limited (herein always referring to lateral diffusion) step, while minimising an unwanted diffusion-limited step. The desired step being that of enzyme-substrate intermediate formation and the unwanted step being that of the intermediate reacting with a second substrate, thereby initiating polymerisation.

A particularly attractive method to overcome the challenges of the conventional bimolecular enzymatic cyclisation reaction would be to make the entire process non-diffusion limited. This can in principle be achieved by fusing the ligase to the substrate (target protein), which would fundamentally alter the cyclisation reaction mechanism from a diffusion-limited bimolecular reaction to a non-diffusion-limited unimolecular reaction. The first order kinetics of the unimolecular reaction allows the sample concentration to be lowered without affecting the reaction half-life. Indeed, elements of this design have appeared in the literature^13, 33, 34^ without consideration or characterisation of the fundamental shift in reaction mechanism that can be achieved, or the gains in cyclisation efficiency that can be realised.

Here, we describe the design of a modular and general “autocyclase” platform for production of head-to-tail macrocyclised proteins and peptides and demonstrate the advantages of first order self-cyclisation (autocyclisation) as a scalable and general protein cyclisation method. We show that under dilute conditions the autocyclases proceed via a unimolecular reaction mechanism following first order reaction kinetics, while suppressing unwanted polymeric side products formed via a bimolecular pathway. The general utility of the autocyclase platform is demonstrated by cyclisation of two popular but very different classes of macromolecules (1) disulfide-rich cyclic peptides and (2) α-helical membrane scaffold proteins (MSPs) for producing circularised nanodiscs (cNDs). We demonstrate the benefits of the facile scale-up of the system by providing the first heteronuclear 3D NMR data of a ^13^C/^15^N isotope-labelled disulfide-rich cyclic peptide. We also produce several nanodisc-based imaging agents and provide some of the first insights into the biodistribution and metabolism of cNDs in vivo. We expect that the versatility of the autocyclases will make these of value in design of novel macrocyclic peptides and proteins as well as improving access to existing cyclic molecules.

## Results

### The autocyclase design

An autocyclase comprises six modules (Figure 1a): (i) a reactive sequence that is liberated by application of a suitable protease (*cleavage site 1*); (ii) a target protein or peptide to be cyclised; (iii) a cyclisation recognition site (*LPGTG)*; (iv) a linker of suitable length and flexibility to promote ligation (*linker*); (v) the cyclising enzyme (*sortase*); and finally (vi) a purification tag (*H10*). An additional module (iv’), with a second (orthogonal) protease cleavage site (*cleavage site 2*), allows the ligase to be isolated as a secondary product (lacking a reactive N-terminal nucleophile—only required if recovery of the enzyme is sought). All autocyclase sequences can be found in the supplementary section (see Supplementary Information and Supplementary Data 1). The two key elements that most significantly affect the efficiency of the autocyclase system are the linker and the ligase, these will be discussed in further detail below.

**Figure 1.**
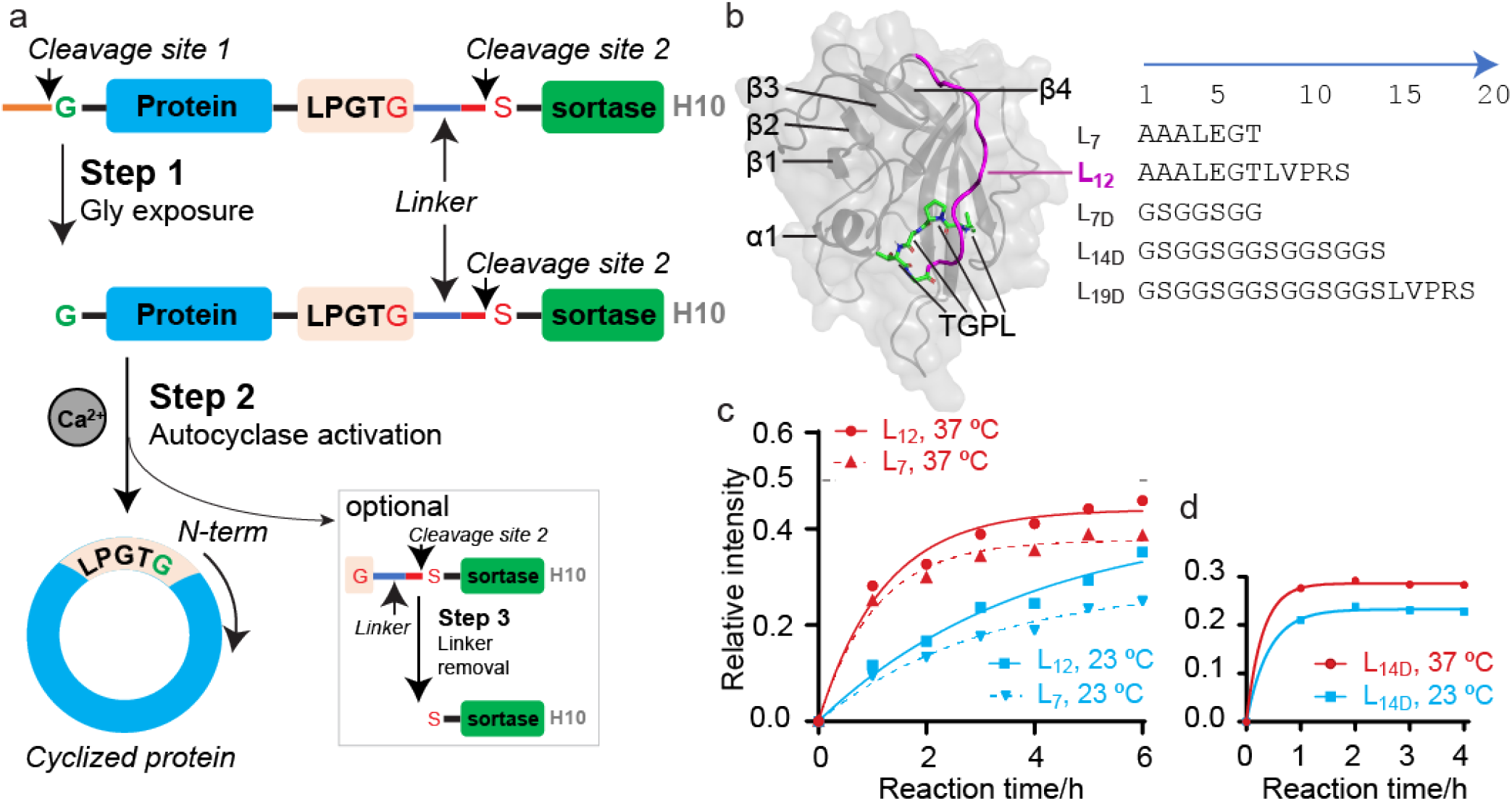
Design of autocyclase as a modular, self-cyclisation platform. **a**, An autocyclase construct features an N-terminal capping sequence (orange line), a protease cleavage site 1 (TEV site— ENLYFQ/G), protein of interest to be cyclised (blue rectangle), SrtA recognition site (pink rectangle), linker (blue line), protease cleavage site 2 (red line, thrombin site—LVPR/S), SrtA (green rectangle) and a C-terminal purification tag (His_10_—H10—in grey). The cyclisation process involves TEV cleavage at cleavage site 1 to expose the N-terminal glycine (step 1), activation of cyclisation reaction using Ca^2+^ ions (step 2) and optionally thrombin cleavage at cleavage site 2 to remove the linker and recover nucleophile-free SrtA (step 3). **b**, Five linkers (L_7_, L_12_, L_7D_, L_14D_ and L_19D_) were designed to connect the protein of interest to the SrtA. MD simulations illustrated that the L_7_ and L_12_ linkers (L_12_ shown in magenta) allow the SrtA recognition site (LPGT residues highlighted in green sticks) to be appropriately positioned at the active site of SrtA. **c-d**, The build-up of cMSP9 from autocyclase (*a*MSP9) cyclisation over 4-6 hours. **c**, The cyclisation rate of cMSP9 was faster for the *a*MSP9-L_12_ than *a*MSP9-L_7_ regardless of temperature (23°C in cyan and 37°C in red). The theoretical and maximum relative intensity of the cyclic product is ∼0.5 (excluding SrtA), indicated by a dash line. The reactions were faster at 37°C for both constructs. **d**, A remarkable increase of reaction rate is observed for the autocyclase containing a dynamic L_14D_ linker.

#### The linker

In our design, a linker connects the SrtA recognition site (LPXTG) to the N-terminal end of SrtA. A suitable linker should allow this recognition site, to access the SrtA catalytic site^35^, across the α1/β2 and β3/β4 loops of SrtA (Figure 1b). The NMR structure (PDBID: 2KID) of SrtA (hereon referring to the Δ59 truncation variant of SrtA*—*residue Q60–K206) reveals that the first secondary structure element (β-strand) starts at residue G74 and that the first 5–9 residues are highly disordered in solution (from ^15^N spin relaxation experiments)^35^. Based on this we estimate that a ∼35– 45 Å, or an approximately 10-amino acid long linker is required (Figure 1b) but note also that some of the disordered N-terminal residues of SrtA may form part of the linker—possibly allowing shorter linkers to be used. A total of five linkers were designed spanning 7–19 residues: 1. [**L**_**7**_ = AAALEGT], 2. [**L**_**12**_ = AAALEGTLVPRS], 3. [**L**_**7D**_ = GS(GGS)GG], 4. [**L**_**14D**_ = GS(GGS)_4_] and 5. [**L**_**19D**_ = GS(GGS)_4_LVPRS]^36^, where the subscript number refers to the number of amino acids and a subscript “D” indicates that the amino acid sequence is designed to yield a dynamic linker.

The production of monomeric cNDs with a 9 nm diameter (cNW9—derived from MSP1D1ΔH4H5 or MSP9) has proven to be particularly challenging^9, 31^, we therefore chose this as a test system to compare the effect of the different linkers on the efficiency of the autocyclase system. Our experiments revealed that autocyclase-MSP9-L_12_ (*a*MSP9-L_12_) produces an increase in quantities of cMSP9 compared to *a*MSP9-L_7_ (Figure 1c, see Supplementary Figure 1 for details). The higher yield of cMSP9 from *a*MSP9-L_12_ compared to *a*MSP9-L_7_ indicates improved yields as a function of linker length. Next, we investigated if introducing residues that would promote disorder in the linker would affect the reaction rate or yields. We designed a linker of similar length to *a*MSP9-L_12_, containing a number of GGS repeats, *a*MSP9-L_14D_, and found a remarkable increase in reaction rate, with the reaction largely complete after ∼1 hour (cf > 6 hours for the L_7_ linkers, Figure 1d). However, we did not see an improvement in overall yields of cMSP9. We also investigated the effect of temperature on the reaction, and found faster reaction rates but with only slightly higher yields of cMSP9 at higher temperatures (Figure 1 and Supplementary Figure 1).

While the longer and more dynamic linkers (L_14D_ and L_19D_) result in a clear increase in reaction rate, this design is also more prone to in vivo hydrolysis leading to a decrease in the overall yields of the *a*MSP9 protein (Supplementary Figure 2). We also introduced a shorter dynamic linker to generate *a*MSP9-L_7D_. Again, we find significant losses due to in vivo hydrolysis, suggesting that linker dynamics alone is sufficient to drive this process (Supplementary Figure 2c). We then introduced an inhibitory peptide (LPRDA)^37^—that binds to the SrtA active site and inhibits the enzyme—at the N-terminal end of the autocyclase (*a-*i-MSP9-L_14D_), but found no improvements (Supplementary Figure 2d), either due to the low affinity of the peptide inhibitor towards SrtA compared to the recognition sequence or due the hydrolysis being driven by other endogenous *E. coli* proteases (e.g. M23 family of bacterial proteases^38^). Thus, our results indicate that the L_12_ linker provides a good compromise between overall reaction rate and overall yield in recombinant production of cyclic MSPs.

Next, we performed molecular dynamics (MD) simulations on the SrtA recognition sequence (LPGTG) linked to SrtA (Q60-K206) directly (L_0_, no linker), but found the complex to be unstable with the recognition sequence leaving the active site within 100 ns (Supplementary Figures 3 and 4). As expected, MD simulations performed with the L_12_ and L_7_ linkers were stable (Supplementary Figure 3); however, significantly different conformations of the linker and the N-terminal SrtA region (Q60-G74) were observed (Supplementary Figure 4). All MD simulations were initiated from the SrtA-substrate structure determined by NMR (PDB ID: 2KID)^35^ which was well reproduced in all three cases (Supplementary Figure 4). The final configurations after 250 ns of MD are included as Supplementary data. Note that simulations of the SrtA-substrate complex performed using the X-ray structure reported by Zong *et al* ^39^ (PDB ID: 1T2W) were unstable, resulting in the substrate leaving the purported active site in all cases.

#### The ligase

The above experiments were all performed using the wild type SrtA sequence (Δ59). We also generated an *a*MSP9-L_7_-eSrtA construct using the evolved sortase pentamutant (eSrtA)^8, 40^ but found it to produce poor overall yields. The low yields were found to be due to (1) enhanced in vivo hydrolysis by the highly reactive enzyme (Supplementary Figure 5a-c) and (2) decreased solubility of the protein when expressed at 37°C (Supplementary Figure 5d-f).

### The autocyclase reaction mechanism

The kinetics and mechanism of the SrtA transpeptidation reaction have been characterised in detail previously under suitable steady-state conditions^41^, while intramolecular enzymatic cyclisation has not previously been studied. In the traditional (bimolecular) enzymatic cyclisation reaction by SrtA, the initial enzyme concentration is typically similar to that of the substrate^8, 9, 31^, and will be expected to display approximately second order reaction kinetics. In the autocyclase (unimolecular) cyclisation reaction (Figure 2a), the initial velocity will be expected to display a mixture of first and second order reaction kinetics (intra- and inter-molecular recognition). Indeed, in the above reactions (using variable linkers) we observe clear evidence of polymerisation at the relatively high concentrations used (∼0.1 mM, see also Supplementary Figure 1), suggesting that the concentration is sufficiently high to allow for intermolecular reactions. However, at lower concentrations, the reaction should favour first order kinetics. It is, therefore, possible to determine a concentration range where the reaction follows a unimolecular mechanism, by following the initial reaction velocity at different (dilute) starting concentration of the autocylcase.

**Figure 2.**
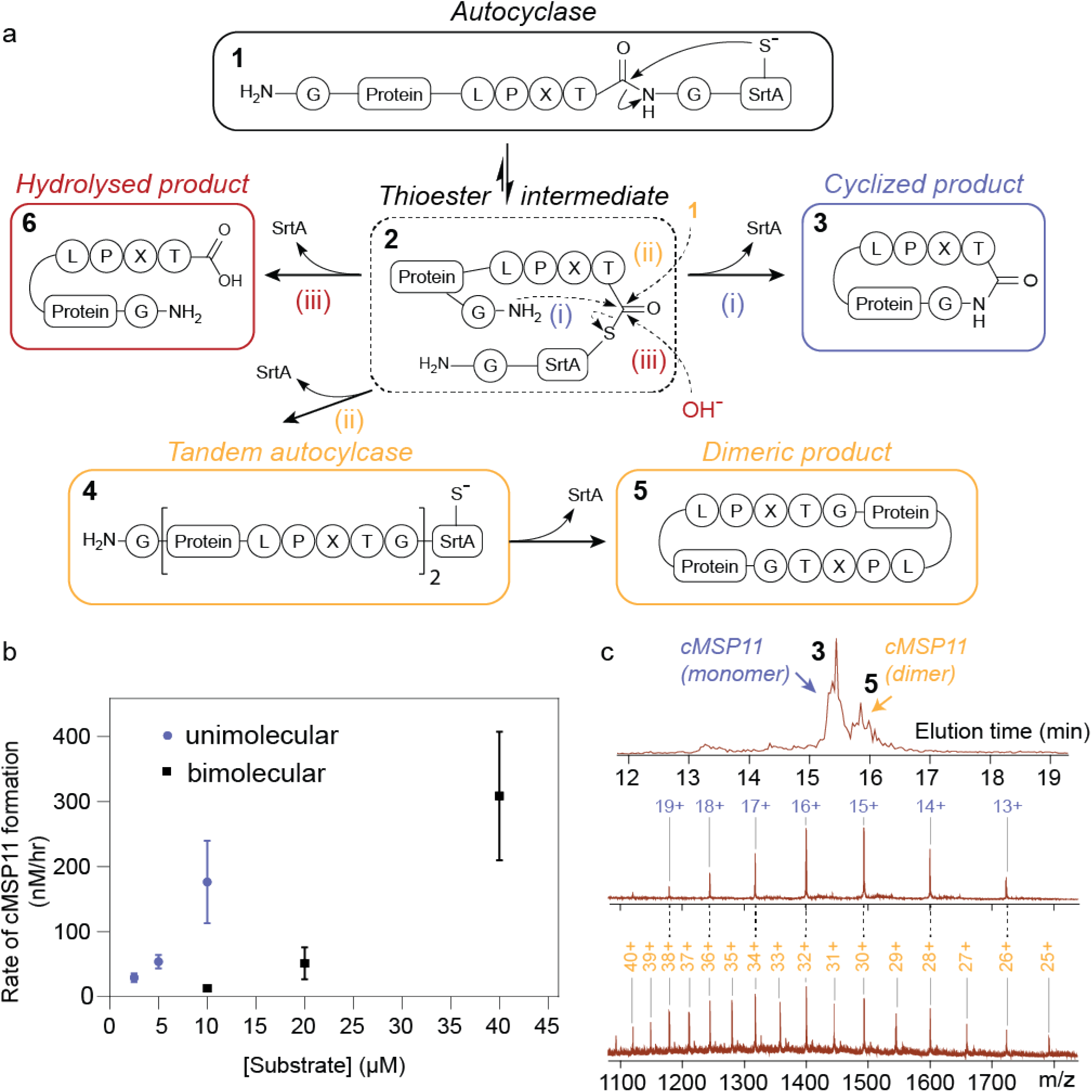
The mechanism of autocyclisation. **a**, A scheme summarising the main reaction pathways of autocyclases. After the generation of a thioester intermediate (2) from a spontaneous intramolecular rearrangement, the reaction can proceed via the following paths: (i) The unimolecular reaction— intramolecular nucleophilic attack by the N-terminal glycine amino group. This pathway results in the cyclised product (3) and free SrtA. (ii) Bimolecular reaction—intermolecular nucleophilic attack. This reaction results in a free SrtA and a protein-protein-SrtA adduct (4—tandem autocyclase). This adduct can self-cyclise to form a cyclic dimer (5). (iii) Hydrolysis—the thioester is resolved by a hydroxide ion and, irreversibly, produces a linear hydrolyzed product (5). **b**, The reaction rates of aMSP11-L_12_ cyclisation are compared to the equivalent reaction using the traditional (bimolecular) enzymatic cyclisation method (using MSP11 and WT SrtA in both cases). Rates were determined by monitoring the build-up of cMSP11 using LC/MS. The rates demonstrate that the autocyclase reaction at lower starting concentrations approximates first order reaction kinetics, while the bimolecular reaction approximates second order reaction kinetics. **c**, The MS spectrum (extracted ion chromatogram) of an aMSP11-L_12_ cyclisation reaction at a high starting autocyclase concentration (200 µM) shows the formation of cyclised monomeric and dimeric products. Identity of the two species were confirmed by the observed m/z ion masses.

Using the *a*MSP11-L_12_ construct we performed a series of reactions at different concentrations at room temperature, and measured the initial rate of cMSP11 formation, by following the reaction progress using liquid chromatography-mass spectrometry (LC/MS—see also Supplementary Information and Supplementary Data 2). The experiment was also repeated under identical conditions using the same MSP11 and SrtA sequences in a traditional enzymatic (bimolecular) ligation reaction. The rates of the autocyclase reaction under dilute concentrations were significantly faster than those of the bimolecular reaction, and while the unimolecular autocyclase reaction was feasible in the range of 2–10 µM, the bimolecular rates were only measurable at or above 10 µM—consistent with the predicted change in reaction mechanism. In the range of 2.5 µm to 5 µM (Figure 2b) we observe a linear change in initial reaction velocity, consistent with first order reaction kinetics (a rate constant of ∼0.01 hr^-1^). Comparing the reaction velocity between 5 and 10 µM, we find a mixture of linear and quadratic behaviour suggesting interference from the bimolecular reaction. In the bimolecular reaction we find that increasing the concentration from 10 µM to 20 µM leads to a quadratic increase in reaction velocity (with a rate constant ∼130 M^-1^hr^-1^), consistent with second order kinetics^41^. Under dilute reaction conditions (∼5 µM), the autocyclase reaction produces the monomeric cyclic product with close to quantitative yields (∼90% of theoretical, Supplementary Tables 1–2).

### Autocyclase by-products

We note that the MS data also indicate that at higher autocyclase starting concentrations (>50 µM) a stable protein-protein-SrtA adduct is formed, which we have termed a “tandem autocyclase” (Figure 2c and Supplementary Figure 6). We find that under suitable conditions, this intermediate, forms a cyclic dimeric product in higher yield than what is produced in the equivalent bimolecular reaction (Figure 2c and Supplementary Figure 6). Depending on the intended application of the cyclised protein, this presents an opportunity to tune the reaction condition for generation of higher order cyclic products.

The MS analysis also revealed the slow build-up of linear MSP over long periods of time. This is the product of the third possible reaction pathway in which the reactive SrtA thioester intermediate is hydrolyzed irreversibly to produce a linear product (see Supplementary Figure 5a)—a common feature of both the autocylase reaction and the traditional bimolecular SrtA reaction.

Finally, we note that some small fraction of unreacted fusion protein often remains upon completion of the reaction. We attribute this to fusion proteins that contain misfolded or otherwise inactivated SrtA species.

### Autocyclases as a general cyclisation strategy

We next investigate the generality of the method, by generating autocyclase constructs containing different sizes of MSPs, as well as several cyclic peptides of pharmaceutical interest. First, utilising the L_12_ linker, we generated cMSP6, 7 and 15 (in addition to cMSP9 and 11 discussed above, see Figure 3 and Supplementary Figures 7–8). Our procedure is very similar to that previously reported^8, 31^, except that we do not need to separately acquire or produce the SrtA enzyme. We also find that the TEV protease cleavage (liberating the N-terminal nucleophile) and cyclisation reactions can be performed concurrently (Supplementary Figure 9), simplifying the process further. We have quantitated our yields at critical steps of the process (Supplementary Tables 1–2), and find that our overall yields are generally higher than those previously reported^8, 9, 31^. We also find that the solubility of our fusion is higher than previous reports of MSPs expressed in *E. coli*, which may be due to the favourable solution properties of the bacterial SrtA protein^8, 9, 31^. We then used the produced cMSP7, 9, 11 and 15 to assemble cNDs (cNW7, 9, 11, 15) containing 1-Palmitoyl-2-Oleoyl-sn-Glycero-3-Phosphocholine (POPC) lipid bilayers (Supplementary Figure 7 and 8). Finally, we used the purified cMSP11 to encapsulate the voltage sensor domain (VSD) of a bacterial voltage gated potassium channel (KvAP)^42^. The ^15^N-TROSY spectrum at elevated temperatures (323 K) shows resonances associated with the correct folding of the protein within the nanodisc (Supplementary Figure 10).

**Figure 3.**
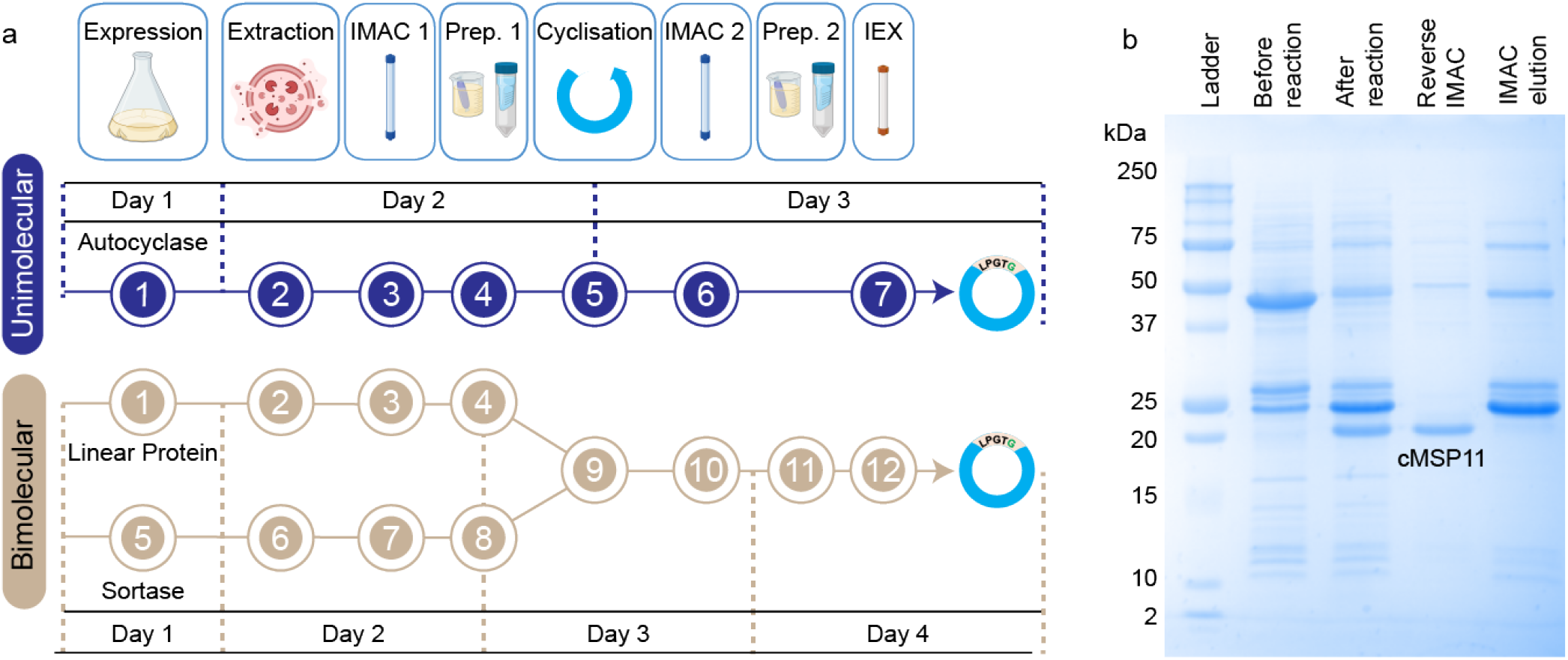
Simplification of enzymatic head-to-tail cyclisation using the unimolecular self-cyclisation method (autocyclisation). **a**, A flowchart outlining each distinct step in the process, comparing autocyclisation with the equivalent bimolecular reaction. Preparation step 1 (Prep. 1) differs between the two methods. In the unimolecular reaction, this step involves dialysis of the fusion protein into a suitable solution for TEV cleavage and cyclisation reactions which are performed as a single step (5). In the bimolecular reaction, this step (4) includes the TEV cleavage reaction as well as a number of buffer exchange steps (preceding and following this step)^8, 31^. **b**, cMSP11 cyclisation using the proposed new method, showing SDS-PAGE lanes corresponding to start and end of steps 5 and 6.

Next, we investigated if the autocyclase method could be employed to produce head-to-tail macrocyclic peptides. We have selected three well-characterised disulfide-rich cyclic peptides that vary in their cysteine content, SFTI (a potent protease inhibitor with one disulfide bond)^43^, Vc1.1 (a potent inhibitor of a nicotinic acetylcholine receptors with two disulfide bonds)^44^ and KalataB1 (kB1; a uterotonic plant peptide with three disulfide bonds)^45^. Initial screening of different linkers in these constructs revealed that these autocyclases were less prone to in vivo hydrolysis compared to the MSPs (Supplementary Figure 2). The longer dynamic linker (L_19D_) could therefore be used to facilitate efficient cyclisation of the cyclic peptides (Figure 4).

**Figure 4.**
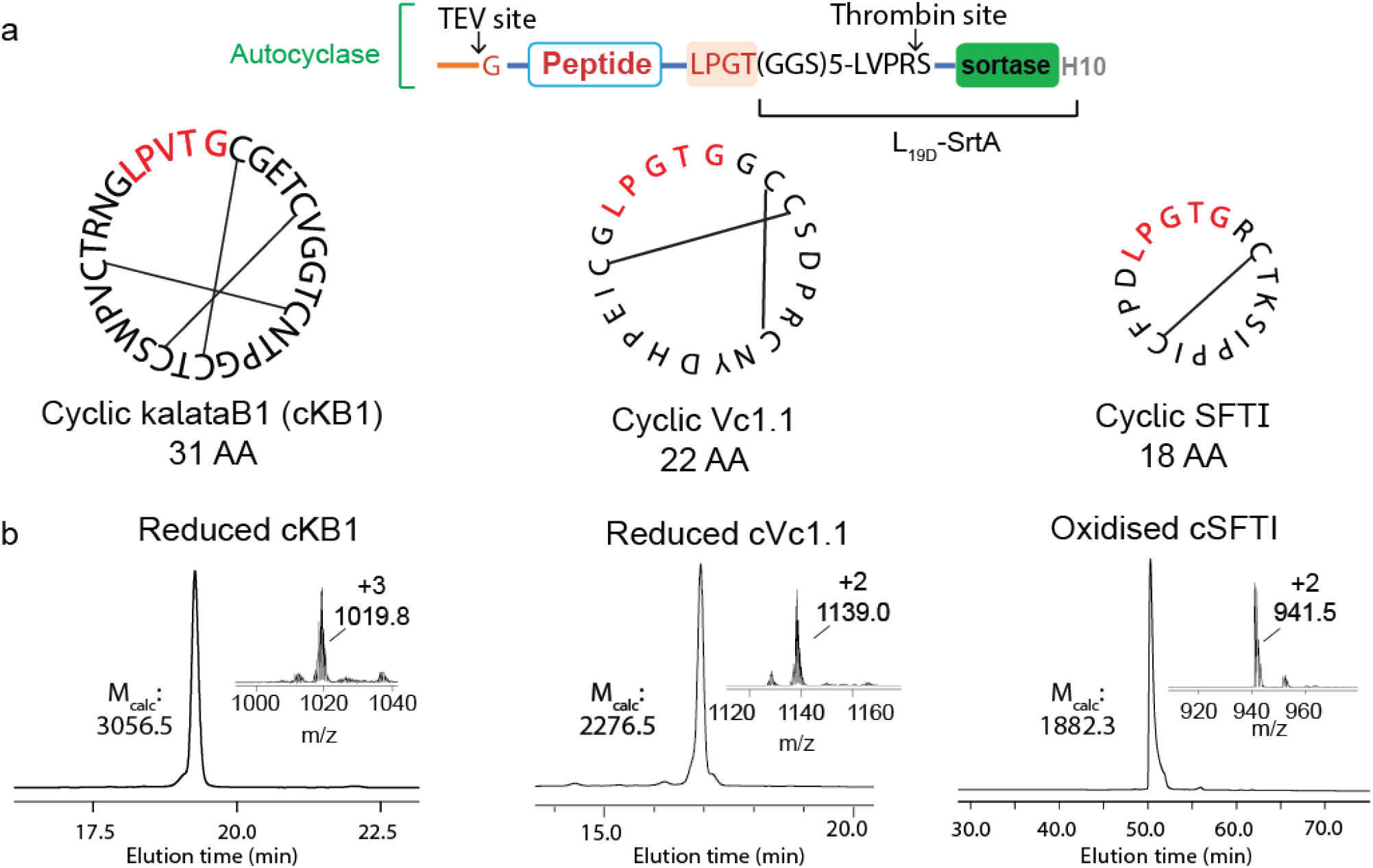
The production of three different cyclic peptides from autocyclases. a, The design of autocyclases for producing cyclic SFTI, kalataB1 (kB1) and Vc1.1. The native peptide sequences are shown in black, with lines indicating the disulfide bond connectivity. The SrtA recognition sequence (LPXTG) that remains in in the peptide after the cyclisation is highlighted in red. **b**, The identity and purity of the cyclised peptides were assessed by rpHPLC and LC/MS. Observed masses were m/z 1019.8 [M+3H]^3+^ for cKB1, m/z 1139.0 [M+2H]^2+^ for cVc1.1 and m/z 941.5 [M+2H]^2+^ for cSFTI. Calculated masses are m/z 1019.8 [M+3H]^3+^ for cKB1, m/z 1139.3 [M+2H]^2+^ for cVc1.1 and m/z 942.2 [M+2H]^2+^ for cSFTI.

Similar to our findings with the MSPs, we found that at higher starting concentrations, there was evidence of the stable tandem autocyclase, yielding cyclised dimers and trimers. In the case of SFTI, we confirmed that as expected the relative proportion of polymeric by-products increases as the starting autocyclase concentration is increased (Supplementary Figure 11).

### Applications of nanodiscs in structural biology and in vivo imaging

The applications of cyclic peptides and nanodiscs are diverse^46^. A number of cyclic peptides have pharmaceutical or agrochemical potentials^7^; while nanodiscs are commonly employed in studies of membrane proteins in a lipide bilayer^8^ and used in imaging and drug delivery applications^47^. Here we demonstrate the utility of the autocyclase system to provide novel insights in both areas.

First, we produced an isotopically-labelled (^13^C/^15^N) SFTI peptide, which allowed us to measure the first triple resonance 3-dimensional (3D) NMR experiments of a macrocyclised disulfide-rich peptide. This peptide sequence had previously been produced using a semi-synthetic approach where the chemically synthesised (linear) peptide was cyclised by SrtA^48^. The produced peptide was observed to yield multiple structural isoforms. The structural heterogeneity was also observed here. While the source of the heterogeneity previously remained unknown, we were able to employ 3D NMR experiments at very high resolution using non-uniform sampling and non-Fourier spectral reconstruction methods^49^, to perform sequential resonance assignment of all structural isoforms of this peptide in solution (Supplementary Figure 12a). We can clearly see 3 isoforms, and using the ^13^C chemical shifts of the proline residues^50^, we are able to conclusively determine that in the major isoform (65% population) all prolines are in the trans configuration, while the two minor isoforms (20% and 15% populations) are due to a cis-proline configuration at positions P13 and P16 respectively (Supplementary Figure 12b)—with the relative energy of formation of each cis-isoform from the predominant trans-isoform calculated to be Δ*G*_*P13*_ = 12 and Δ*G*_*P16*_ = 15 kcal·mol^-1^. Furthermore, we are able to unequivocally demonstrate the presence of a covalent bond between T18 and G1 in all three isoforms (Supplementary Figure 12c).

Another important application of nanodiscs, is their use as carriers of drugs or imaging agents^47^—not least in the form of peptides^7, 45, 51, 52^. Despite their potential utility in this area little is known about how the size or cyclisation of nanodiscs affect their in vivo fate (in absence of a cargo)^47, 53^. To probe this, we generated both linear and cyclised nanodiscs (NW11 and cNW11) carrying two optical probes that fluoresce at different wavelengths—one inserted into the lipid bilayer, and the other conjugated to the MSP (Figure 5). We also generated small (cNW7) and large (cNW15) diameter dual-labelled cNDs. The two sets of nanodiscs allow us to monitor the effect of cyclisation and size on biodistribution and metabolism in vivo. Indeed, we find clear differences in both biodistribution and metabolism across the different samples over the 24 h post intravenous injection (Figure 5, Supplementary Figure 13, Supplementary Data 3).

**Figure 5.**
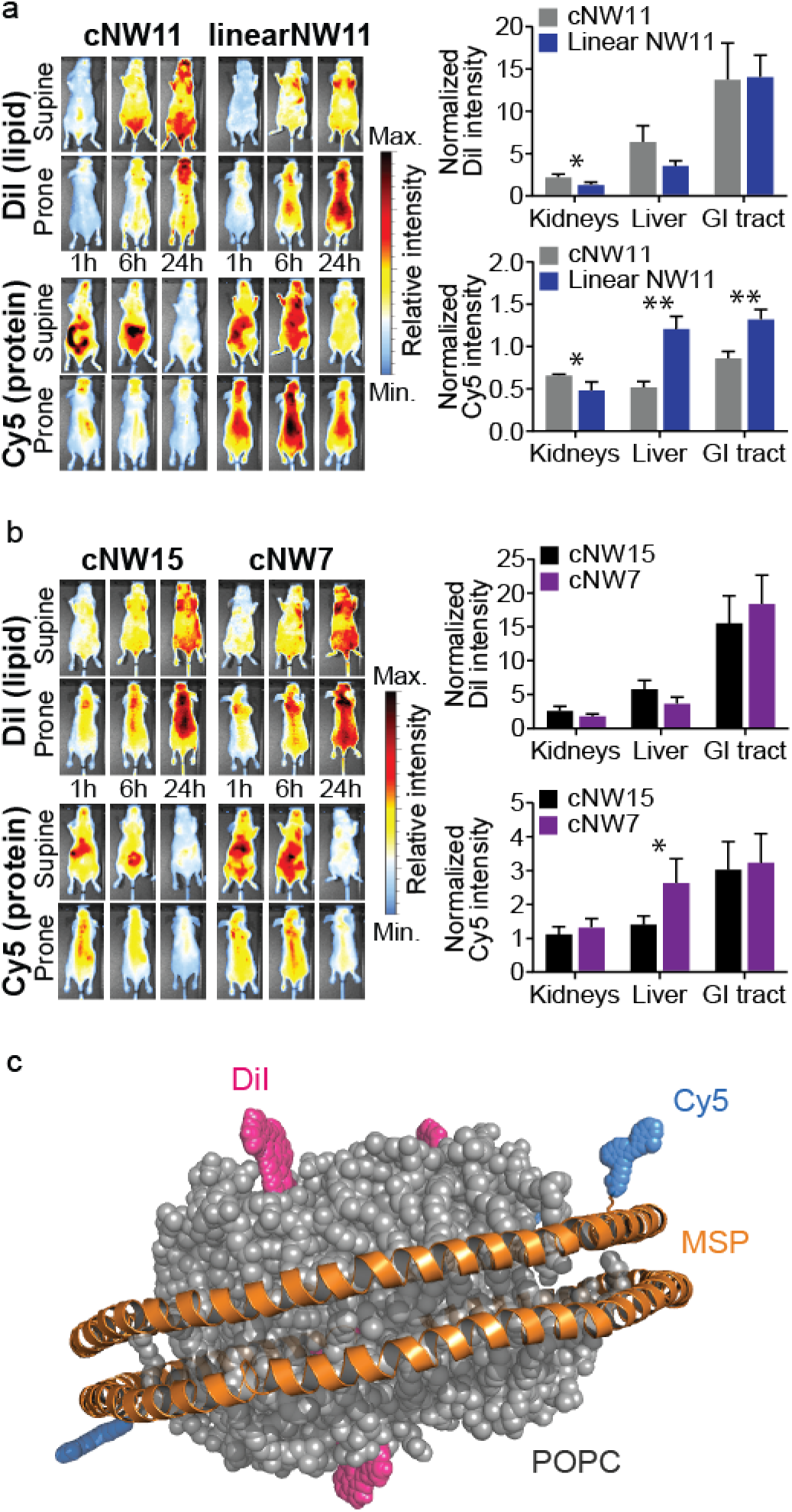
In vivo biodistribution of nanodiscs in naïve BALB/c nude mice. Animals were administered fluorescent nanodiscs via the tail vein and imaged 1, 6, and 24 hours post injection. Imaging of the Cy5 conjugated MSP protein was performed using 620 nm excitation and 670 nm emission filters and the lipophilic DiI dye using 540 nm excitation and 620 nm emission filters. **a**, Comparison of cNW11 and linear NW11 distributions. DiI intensity, localised to regions of fat deposits, increases with time reaching a peak at 24 hours post injection for both linear and cyclised nanodiscs. Cy5 distribution demonstrates clearance of both linear and cyclic nanodiscs through the GI tract at early time points. Ex vivo analysis of clearance organs demonstrates higher linear NW11 protein in the liver 24 hours post injection compared to cNW11 but lower DiI suggesting increased metabolism. Ex vivo organ fluorescence data is presented as mean normalised radiant efficiency values (n=3) with standard deviation error bars. **b**, Comparison of cNW15 and cNW7 distributions. Both cNW15 and cNW7 demonstrate similar distributions to cNW11. Ex vivo analysis demonstrates higher cNW7 protein in the liver compared to cNW15, similar to the trend observed between linear NW11 and cNW11. Statical analysis was performed using unpaired t-tests, p-value ≤ 0.05 (*), ≤ 0.005 (**). **c**, Graphical representation of a fluorescent nanodisc demonstrating the Cy5 (blue) conjugated to the MSP (orange) lysine sidechain, and the lipophilic DiI dye (magenta) located within the POPC lipid bilayer (grey).

The lipid content of the discs appears to preferentially accumulate in brown adipose tissue (see fat pads near the animal head), while the MSPs are generally found in the regions associated with clearance (liver, kidney and GI tract) (Figure 5). Comparing the linear and cyclic proteins (Figure 5a), we find higher levels of the linear MSP in the liver (p value 0.002) and GI tract (p value 0.005) compared to cyclic MSP (normalised to blood), supporting increased in vivo stability following cyclisation. Similarly, the cND appears to deposit more lipids in the liver compared to the linear nanodiscs, consistent with a longer circulation time. Comparison of the *in vivo* images of smaller and larger cNDs (cNW7 and cNW15; Figure 5b), showed the smaller discs to be cleared more rapidly than the larger discs, with similar patterns in clearance organ accumulation at 24 h post administration. The only statically significant difference observed between small and large cNDs was an increase in the smaller cMSP protein in the liver (*p*-value 0.05), associated with increased clearance. These data provide the first insights into the in vivo properties of cNDs and suggest that larger cNDs are more resistant to clearance than smaller and linear nanodiscs.

## Discussion

Here, we describe the engineering of a class of proteins, termed autocyclases, that can undergo intramolecular head-to-tail macrocyclisation to release an enzyme and a cyclic peptide or protein. We show that conversion of the autocyclase to a monomeric cyclic product, increases as a function of decreasing starting concentration, while approximating first-order reaction kinetics, reaching nearly quantitative yield below concentrations of ∼5 µM. We find that under identical conditions the equivalent bimolecular reaction is unfeasible.

In recent years, several methods have emerged that enable in vivo production of self-cyclised MSPs^34, 54^. This is achieved by flanking the MSP sequence with two reactive elements that upon expression can spontaneously yield a cyclic product. The first of these, uses the intein-mediated ligation method to achieve head-to-tail cyclisation^54^, while the other uses the SpyCatcher/SpyTag technology to produce a head-to-sidechain cyclised product.^34^. While the former leaves a modest ligation scar the latter yields a product carrying a ∼120 amino acid insertion. Both methods typically also require the inclusion of an affinity tag within the cyclised product for downstream purification. In vivo cyclisation makes these methods attractive as they dramatically improve throughput. In vivo cyclisation, however, also prohibits detailed characterisation and control of the reaction. The main difference of these methods to that present here is that the cyclisation reaction is here triggered in vitro, upon cleavage of the protecting N-terminal capping sequence and addition of calcium. As demonstrated, we can use this property to characterise the reaction in detail, providing opportunities to tune the reaction conditions or explore how small changes to the protein sequence (enzyme or linker) can affect the outcome of the cyclisation reaction—allowing for further optimisation of the self-cyclisation reaction.

In the autocyclase reaction, the key departure from traditional (bimolecular) enzymatic cyclisation is the first step of the reaction (Figure 2a), i.e. the recognition of the substrate sequence by the enzyme. In the bimolecular SrtA reaction, this is described by the *K*_m_ of the enzyme, i.e. directly related to the substrate concentration. In the unimolecular reaction (using the same enzyme and recognition motif), this step is concentration independent and relates instead to the properties of the linker (frequency and orientation of collisions of recognition sequence with the catalytic site). Once the intermediate is formed, all subsequent steps are largely the same – further emphasising the significance of the linker.

Interestingly, several class A and class C sortase enzymes contain a flexible, N-terminal segment that may regulate substrate binding because their structures reveal that this N-terminal appendage partially shields the active site and wraps around the surface of the catalytic site^55^. It is likely that our design is hijacking this natural autoregulatory function, by replacing the autoinhibitory sequence—that has evolved to access the catalytic site—with a recognition site for autocyclisation. This is supported by our MD simulations, which further show that direct fusion of the target protein to SrtA lacking an N-terminal spacer, leads to an unstable complex (Supplementary Figures 3 and 4). Interestingly, we note that in a reported example of human growth hormone (hGH) cyclisation,^33^ a direct fusion of hGH to SrtA was created to enable “one-step” purification and cyclisation of hGH. The possibility of intramolecular cyclisation was not considered, and given the results of our MD simulations we can conclude that the system although reducing the number steps of SrtA mediated cyclisation (see Figure 3), achieved cyclisation via the traditional intermolecular reaction.

While characterising the unique properties of protein self-cyclisation we have only performed limited optimisation of the system. An important improvement would be to engineer the system to be resistant to polymerisation at concentrations higher than what is currently demonstrated (∼5 µM). As noted above, the difference between intramolecular cyclisation and intermolecular polymerisation will depend on the difference between the frequency of successful collisions of the enzyme active site and the recognition sequence—the former driven by linker dynamics and the latter by substrate concentration. Above a critical concentration, determined by *K*_*m*_ of LPXTG recognition, the quadratic increase in the intermolecular reaction rate will rapidly make this the dominant reaction path. The *K*_*m*_ of WT-SrtA has been reported to be ∼10-100 µM^35, 41^, and indeed we find that in this range (>10 µM), the bimolecular reaction significantly interferes with unimolecular cyclisation, setting an upper concentration limit on the utility of the unimolecular path. However, we would predict that an increase in the *K*_*m*_ would increase the critical concentration where we see oligomerisation of the substrate. Mutations that affect the SrtA recognition site are known to affect the *K*_*m*_, increasing it up to 6-fold^35^. It is therefore possible that such a mutation may allow for the upper limit of the unimolecular reaction to be increased. Combining such a mutation with an optimised linker would be particularly interesting, as any losses in reaction rate due to the weaker binding of the recognition sequence to the recognition site, would be compensated by the increased effective local concentration at this site due to favourable linker dynamics. Such optimisation can now be pursued following the establishment of the described theoretical framework. Autocyclases, thus, offer a flexible system that through continued development promises to lead to further advances in enzymatic head-to-tail macrocyclization.

Finally, we demonstrate the utility of macrocyclised proteins produced using the autocyclase approach by (1) uniformly ^13^C/^15^N isotope-labelling a cyclised disulfide-rich peptide for the first time, (2) generating a range of cNDs of different sizes, including one containing an ion channel voltage sensor domain, and (3) providing the first data on the in vivo biodistribution of cNDs and clearance pathway as a function of cND size and cyclisation. Isotope labelling demonstrate the wealth of structural NMR data which can now be generated through isotope labelling of cyclic peptides, while our imaging experiments provide evidence that large cNDs may serve as useful carriers of drugs and imaging agents.

Clearly the applications of macrocyclised peptides and proteins are both numerous and diverse. Access to a simple, fast and low-cost method of producing these will facilitate future research across fields of structural biology and pharmaceutical sciences.

## Conclusion

Cyclic peptides and proteins have many properties that make them attractive as biochemical tools and for pharmaceutical development. Numerous enzymatic methods have emerged that are capable of cyclising linear substrates. Here we present a method that incorporates both the enzyme and the substrate in the same molecular entity. We show that this leads to a change in reaction mechanism, resulting in a self-cyclisation reaction following first-order reaction kinetics. This change from a diffusion limited path to a non-diffusion limited one, presents new opportunities to optimise reaction conditions that favour cyclisation over polymerisation. The utility of the method is demonstrated by production and characterisation of a range of cyclic peptides and proteins.

## Experimental Section

### Cloning

All sequences generated in this study and the primers used in their production are provided in the Supplemental sections (Supplementary Tables 3–4 and Supplemental Data 1). All autocyclase constructs described here feature an N-terminal capping sequence, a TEV cleavage site, the MSP or peptide of interest, a linker of various lengths and sequences, SrtA (evolved or WT) and a C-terminal His_10_ tag. As a template, a codon-optimised gene encoding an N-terminal His_6_, TEV site, MSP9, L_7_ linker, eSrtA and a C-terminal His_6_ was purchased from IDT. The gene was double digested with *Nde*I and *Xho*I, cleaned up using a macherey-nagel nucleospin gel and PCR clean-up kit and cloned into a pET29a vector. Genetic modifications on this template plasmid were achieved by either gene mutagenesis using NEB Q5 mutagenesis kit or gene replacement using restriction enzyme digestion, to obtain other constructs that encode various MSP, linkers, sortase or histidine tags. The amino acid sequence of the MSPs in these constructs are based on previous reports (including MSP9 and MSP11^8^, MSP6 and MSP7^56^, and MSP15^57^). The macrocyclic peptide autocyclase constructs were produced by replacing the MSP9 gene in the *a*MSP9 autocyclase template. The peptide sequences were selected based on past literature reports^43-45^. Detailed methods describing specific cloning protocols for each construct are provided in the Supplementary Information.

### Autocyclase expression and purification

Each autocyclase expression construct (in a pET29a vector) was transformed into *E. coli* BL21(DE3) cells. Freshly transformed colonies or glycerol stock were used to inoculate LB media containing 50 µg/mL of kanamycin. The starter culture was incubated at 30°C and agitated (shaking at 220 rpm) overnight. 3 mL of the preculture was used to inoculate 300 mL of LB broth containing 50 µg/mL of kanamycin. The culture was then incubated at 37°C and agitated (shaking at 250 rpm) until the OD_600_ reached ∼1.0. Expression was induced by addition of 0.2 or 1 mM IPTG and the culture was left shaking at 250 rpm for 1–6 h at 30°C. The cells were harvested at 6 h by centrifugation at 6,000 *g* for 10 min at 4°C.

The cell pellets were resuspended in lysis buffer (25 mM sodium phosphate pH 7.4, 500 mM NaCl, 20 mM imidazole) containing 1 mg/mL lysozyme and stirred at 4°C for 0.5 h. The resuspended cells were lysed by two 5-minute cycles of sonication on ice (digital sonifier 450 Branson; 40% power; repetitions of 3 s on-pulse and 12 s off-pulse) with a 5-minute break between the cycles to avoid overheating. The sonicated sample was centrifuged at 30,000 *g* for 30 min at 4°C to remove the insoluble fractions and the supernatant was loaded onto a gravity column containing Ni-NTA resin (pre-equilibrated with 4°C lysis buffer). The resin was washed with five column volumes (CVs) of lysis buffer and the autocyclase was eluted with five CVs of elution buffer (25 mM sodium phosphate pH 7.4, 500 mM NaCl, 500 mM imidazole).

### MSP and peptide cyclisation

The cyclisation process follows the steps described in Figure 3. For quantitation of the cyclisation reaction, the first preparation step (4) includes an anion exchange chromatography step to remove any free sortase enzyme that is co-purified with the fusion protein. While this affects quantitation required to determine the reaction mechanism, it can be excluded to make the process more efficient without significantly affecting the final yields. Details of each step in Figure 3 are provided in the Supplementary Information.

Further details on the reagents, cloning, over-expression, purification, cyclisation, SDS-PAGE analysis, enzyme kinetics, Molecular dyanmics, empty nanodisc and KvAPVSD nanodisc asembly, NMR experiments and biodistribution analysis were described in the Supplementary Information.

## Acknowledgements

This work was supported by the Australian Research Council (DP220103028 to MM and LP180100486, CE140100036 and IC170100035 to KJT), the National Health and Medical Research Council of Australia (APP1162597 to MM and APP1148582 to KJT) and the University of Queensland (Research Stimulus fellowship to YC and Development Fellowship to MM). The authors are indebted to Prof. Gerhard Wagner and Assoc. Prof. Mahmoud Nasr for hosting XJ. at Harvard Medical School and providing valuable training in nanodisc experiments. We thank Prof. Zakhar Shenkarev for providing NMR data and advice on KvAP nanodiscs assembly. The plasmid for expressing evolved SrtA pentamutant (P94R/D160N/D165A/K190E/K196T; eSrtA) was kindly provided by Prof. David Liu at Harvard University. The authors are grateful to the Queensland NMR Network and the Centre for Microscopy and Microanalysis at The University of Queensland (UQ) for providing training, access and support in use of NMR and transmission electron microscopy instruments.

