## supplementary information for "Self-cyclisation as a general and efficient platform for peptide and protein macrocyclisation"

#### Supplementary Methods

*Reagents*

All lipids were purchased from Avanti Polar Lipids, Inc. (Alabaster, AL) or Sigma-Aldrich. Enzymes and buffers used for polymerase chain reactions (PCR) and molecular cloning were purchased from Genesearch, the exclusive Australian distributor of New England Biolabs (NEB) molecular biology products. The Quick-stick ligase was purchased from Bioline. The DNA miniprep kit was bought from Qiagene. Gel extraction and PCR clean-up kit were purchased from Macherey-Nagel. All Sanger Sequencing services were acquired from Australian Genome Research Facility (AGRF). All primers and codon-optimised gene fragment were order from Integrated DNA technology (IDT).

*The molecular cloning of autocyclase for MSPs*

The amino acid sequences of the MSPs generated here (see Supplementary Data 1) were based on literature reports (including MSP1D1ΔH5 or MSP9 and MSP1D1 or MSP11[1], MSP1D1ΔH4-6 or MSP6 and MSP1D1ΔH4-5MSP7[2], and MSP2N2 or MSP15[3]). A codon-optimised gene fragment was designed to encode: N-terminal $His_6$, TEV site, MSP9, linker, evolved sortase A (eSrtA) and a C-terminal $His_6$ ($His_6$-TEV-MSP9-LPGTG-$L_7$-eSrtA-$His_6$). This sequence also includes a *Kpn*I site (GGTACC) between the $L_7$ linker and the eSrtA sequence (used later). This gene was ordered from IDT and digested with *Nde*I/*Xho*I to generate an overhang for cloning into the pET29 vector, and subsequently purified using a PCR clean up kit. An expression plasmid which encodes eSrtA in a pET29a vector was then digested by *Nde*I/*Xho*I[4] and purified by DNA agarose electrophoresis. The digested $His_6$-TEV-MSP9-LPGTG-$L_7$-eSrtA-$His_6$ gene was then ligated into the digested pET29a vector.

To generate the fusion protein $His_6$-TEV-MSP9-LPGTG-$L_7$-(WT)SrtA-$His_6$, wild type SrtA-staph-Δ59 (henceforth SrtA) was amplified using the P1 primer pair (Supplementary Table 3). The PCR product was digested with *Kpn*I and *Xho*I and used to replace the eSrtA fragment in $His_6$-TEV-MSP9-LPGTG -$L_7$-eSrtA-$His_6$ to yield $His_6$-TEV-MSP9-LPGTG-$L_7$-SrtA-$His_6$.

The removal of the N-terminal $His_6$-tag in $His_6$-TEV-MSP9-LPGTG-$L_7$-SrtA/eSrtA-$His_6$ was achieved using the P2 primer pair while the introduction of four more histidines into the $His_6$-

tag at the C-terminus was achieved using the P3 primer pair (Supplementary Table 1), yielding TEV-MSP9-LPGTG-L$_{7}$-SrtA/eSrtA-His$_{10}$.

The linker replacement in TEV-MSP9-LPGTG-L$_{7}$-SrtA/eSrtA-His$_{10}$ was achieved using the primer P4 pairs and P5 pairs respectively (Supplementary Table 1), yielding TEV-MSP9-LPGTG-L$_{14D}$-SrtA/eSrtA-His$_{10}$.

To generate the fusion of TEV-MSP11-LPGTG-L$_{14D}$-SrtA-His$_{10}$, MSP11 was amplified from MSP1D1[1] expression plasmid using the P6 primer pair. The PCR product was digested with *Nde*I/*Kpn*I to replace the MSP9 fragment to TEV-MSP11-LPGTG-L$_{14D}$-SrtA-His$_{10}$.

To generate the TEV-MSP15-LPGTG-L$_{14D}$-SrtA-His$_{10}$ fusion, a codon optimised MSP15 gene block (IDT) was digested with *Nde*I/*Kpn*I to replace the MSP9 fragment in the TEV-MSP9-LPGTG-L$_{14D}$-SrtA-His$_{10}$ fusion to yield TEV-MSP15-LPGTG-L$_{14D}$-SrtA-His$_{10}$. Primer pair P7 was used to delete Helix4 in the MSP9 construct to generate TEV-MSP7-LPGTG-L$_{14D}$-SrtA-His$_{10}$. Primer pair P8 was used to delete Helix4 and Helix6 in the MSP9 construct to generate the corresponding TEV-MSP6-LPGTG-L$_{14D}$-SrtA-His$_{10}$ (Supplementary Data). Primer pair P9 was used to introduce a SrtA inhibitory peptide (*i*) for the potential in vivo inhibition of SrtA activity to generate *i*-TEV-MSP9-LPGTG-L$_{14D}$-SrtA-His$_{10}$. Primer pair P10 was used to introduce a thrombin site between the L$_{7}$-linker and SrtA while P11 was used to do the same between the L$_{14D}$ linker and SrtA – yielding TEV-MSP9-LPGTG-L$_{12}$-SrtA-His$_{10}$ and TEV-MSP9-LPGTG-L$_{19D}$-SrtA-His$_{10}$. P7 was then used to generate TEV-MSP7-LPGTG-L$_{12}$-SrtA-His$_{10}$ and P8 to generate TEV-MSP6-LPGTG-L$_{12}$-SrtA-His$_{10}$. For producing TEV-MSP11-LPGTG-L$_{12}$-SrtA-His$_{10}$, a gene encoding His$_{6}$-MSP11 was cut out from a plasmid containing the MSP11 gene with *Nde*I/*Kpn*I, and was subsequently ligated into the vector containing TEV-MSP9-LPGTG-L$_{12}$-SrtA-His$_{10}$ from which MSP9 had been removed. Finally, the N-terminal his-tag of MSP11 was removed by the primer pair P12. For constructing TEV-MSP15-LPGTG-L$_{12}$-SrtA-His$_{10}$, the MSP15 gene (in a pUCIDT-AMP+ vector (IDT)) was obtained and primer pair P13 was then used to amplify MSP15, which was then digested with *Nde*I/*Kpn*I and used to replace MSP9 in TEV-MSP9-LPGTG-L$_{12}$-SrtA-His$_{10}$ (Supplementary Data).

*The molecular cloning of autocyclase for cyclic peptides*

We use the primer pair P14 to delete MSP9 in *i*-TEV-MSP9-LPGTG-$L_{19D}$-SrtA-$His_{10}$ to generate an empty autocyclase vector (*i*-TEV---LPGTG-$L_{19D}$-SrtA-$His_{10}$). Primer pairs P15 and P16 were then used to inserted G-SFTI and G-kB1 respectively between the TEV-site and the $L_{19D}$ linker, in the empty vector. To generate G-SFTI and G-kB1 in an autocyclase with an $L_{12}$ linker, we ordered a gene block encoding kB1 (IDT), digested with *Nde*I/*Kpn*I and replaced MSP9 and LPGTG in TEV-MSP9-LPGTG-$L_{12}$-SrtA-$His_{10}$ to generate TEV-G-kB1-LPVTG-$L_{12}$-SrtA-$His_{10}$.

TEV-GG-Vc1.1-LPGTG-$L_{12}$-SrtA-$His_{10}$ was constructed by replacing MSP9 in TEV-MSP9-LPGTG-$L_{12}$-SrtA-$His_{10}$ with GG-Vc1.1 using primer pair P18. Primer pair P19 was then used in PCR mutagenesis to replace $L_{12}$ with $L_{19D}$ to produce TEV-GG-Vc1.1-LPGTG-$L_{19D}$-SrtA-$His_{10}$.

*Cyclisation of MSPs and peptides*

To expose the N-terminal glycine, the autocyclase in cleavage buffer (25 mM Tris·HCl pH 7.5, 150 mM NaCl, 1 mM β-mercaptoethanol (BME), 0.5 mM EDTA) was cleaved by TEV protease at a TEV-to-protein ratio of 1:50 (w/w) at 4°C overnight. The cleaved protein sample was then spun down at 3000 *g* for 5 min at 4°C to remove any precipitates.

To acquire MSP-autocyclase samples with high purity (for kinetic and cyclisation-efficiency studies), the TEV-cleaved sample was further purified using anion exchange chromatography. The sample was dialysed into buffer containing 25 mM Tris·HCl pH 7.5 and applied to a HiTrap Q Fast Flow column (3 mL; Cytiva). The autocyclase was eluted with 5 CVs of buffer containing 25 mM Tris·HCl pH 7.5 and 50 mM NaCl and was dialyzed into the reaction buffer (25 mM Tris·HCl pH 7.5, 150 mM NaCl) for cyclisation.

For MSP cyclisation, the autocyclase in the reaction buffer was supplemented with 1 mM BME, 10 mM $CaCl_2$ was added to initiate the reaction. This reaction can be performed together with the TEV cleavage reaction to improve throughput, without affecting outcomes. Various conditions were tested in this study to optimise the yield of the cyclic products. The variables include (i) initial autocyclase concentrations (5-100 µM); (ii) with and without the supplement of detergents (1-2 mM Triton X-100 or DDM); (iii) reaction temperatures (23°C or 37°C), and (iv) reaction time (1-

18 h). The formation of reaction products was monitored and assessed using SDS-PAGE or liquid chromatography-mass spectrometry (LC-MS).

For SFTI, Vc1.1 and kalata B1 cyclisation, the samples were dialyzed into the reaction buffer after TEV cleavage. The reactions were initiated by the addition of 10 mM $CaCl_2$ and carried out at 37°C overnight. The reactions were conducted at 50, 100 or 200 µM autocyclase concentration during optimisation. For large-scale production, the concentration was kept below 100 µM. For SFTI, the reaction was supplemented with 3 mM reduced and 0.3 mM oxidised glutathione to produce oxidised cyclic peptides. For Vc1.1 and kalata B1, 3 mM BME was used in the reaction mixture to produce reduced peptides. The reaction completeness and yield were assessed by LC-MS.

To purify the cyclic products, the reaction mixture was passed through Ni-NTA resin on a gravity column that was pre-equilibrated with the reaction buffer (25 mM Tris·HCl pH 7.5, 150 mM NaCl). The flow-through containing the untagged cyclic product was collected.

*Determination of aMSP cyclisation efficiency*

Cyclic MSPs (produced as above at 5 µM starting concentration at 37°C for 18 h) were diluted to 1 µM and then analyzed by reverse phase high performance liquid chromatography (rpHPLC) using an analytical C18 column (Agilent) to determine the ratio of cyclic monomer to dimer. The HPLC gradient was either from 40–60% or 40%-80% (of solvent B) at a slope of 2% per min, at a flow rate of 1 mL/min (solvent A: water with 0.05% TFA; solvent B: 90% $CH_3CN$, 9.957% water, and 0.043% TFA). The ratio of cyclic monomer to dimer was calculated by the area of each peak based on the HPLC profile. The conversion rates were calculated based on the $A_{280}$ of the reverse $Ni^{2+}$-IMAC flow-through, subtracting impurities following HPLC analysis.

*Cyclic MSP purification*

For the cMSP, the Ni-NTA flowthrough fractions were dialyzed into equilibration buffer (25 mM Tris·HCl pH 8, 1 mM DDM) using an Amicon 10 kDa molecular weight cut-off centrifugal filter (Merck). The sample was then purified using anion-exchange chromatography with a HiScreen™ Capto™ Q column (Cytiva) on an ÄKTA Purifier FPLC system (GE Healthcare) at 4°C. The flow rate was 0.3 mL/min and a linear gradient from 0 to 20 % of equilibration buffer supplemented with 1 M NaCl was applied over 8 CVs. Chromatograms monitoring the $A_{280}$ were recorded, and 4 mL fractions were collected by an automated fraction collector (Frac-920) throughout the run. Fractions containing cMSPs with purity >95% as judged by SDS-PAGE were pooled, flash-frozen for storage or used directly for nanodisc assembly.

*Cyclic peptide purification*

Ni-NTA flow-through fractions containing the cyclic peptides were acidified with 1% trifluoroacetic acid (TFA) and filtered before being purified by rpHPLC. Peptide samples were loaded directly onto a semi-prep C3 column (Agilent Zorbax® 300SB-C3 5 µm, 9.4 x 250 mm) using a HPLC system (Agilent Technologies 1260 Infinity). The column was equilibrated with 90% of solvent A ($H_2O$:TFA 99.95:0.05) and 10% solvent B (acetonitrile (ACN):$H_2O$:TFA 79.95:20:0.05) and the chromatography was performed at a flow rate of 3 mL/min with a linear gradient of 20–80% of solvent B over 30 min. Fractions containing peptides were pooled and further purified on an analytical C18 column (Agilent Zorbax® 300SB-C18 5 µm, 4.6 x 250 mm) with a linear gradient of 20–80% of solvent B over 30 min. Sample identity and purity were assessed by MALDI-TOF/TOF MS (Bruker Daltonics Autoflex Speed). Fractions containing the pure peptides were pooled and lyophilised for storage.

*Sortase A regeneration*

After the autocyclase cyclisation reaction, free SrtA containing an N-terminal linker (as well as any unreacted autocyclase) is captured on the Ni-NTA resin in the reverse IMAC purification step. If required, it is possible to capture this enzyme and treat it for future use in other applications. For this purpose the resin containing the capture linker-SrtA fusion was rinsed with 5 CVs of thrombin buffer (20 mM Tris·HCl pH 7.5, 50 mM NaCl) and incubated with thrombin protease (Sigma-Aldrich) at a thrombin-to-‘linker-SrtA’ ratio of 1 unit : 1 mg at room temperature for 6 h or overnight. After thrombin cleavage, the resin was washed with 5 CVs of the thrombin buffer and the regenerated SrtA (lacking the linker) was eluted with the SrtA elution buffer (25 mM sodium phosphate pH 7.4, 500 mM NaCl, 500 mM imidazole). Pure SrtA was buffer exchanged into SrtA buffer (20 mM Tris·HCl pH 7.5, 100 mM NaCl, 2 mM DTT) by an Amicon 10 kDa molecular weight cut-off centrifugal filter (Merck) for storage at -80°C.

*SDS-PAGE analysis*

Electrophoresis on all protein samples was conducted using homemade SDS-PAGE gels (12 or 15% acrylamide) in tris-glycine running buffer at 180–220 V for 30–50 min. Protein samples were prepared by mixing equal volumes of protein sample and 2x SDS-PAGE loading dye and heating at 95 °C for 10 minutes. An extra step of vortex to reduce sample viscosity was introduced for preparing whole cell lysate samples. The Precision Plus Protein Dual Xtra ladder (Bio-Rad) was used as molecular weight standard markers. After electrophoresis, gels were thoroughly washed with hot distilled water before staining with Coomassie Brilliant Blue (CBB) and destained with distilled water[5]. Band intensity of Coomassie staining was quantified using ImageLab software (Bio-Rad). Cyclisation efficiency was calculated as the intensity ratio of the cyclic product band divided by the intensity of starting material (autocyclase) band.

*Enzyme kinetics*

We compared the rates of MSP11 cyclisation by monitoring an autocyclase (*a*MSP11-$L_{12}$) reaction and the equivalent MSP11 and free WT SrtA reaction. The kinetics were quantitated using LC-MS (Thermo Fisher UltiMate 3000 HPLC interfaced with Bruker micrOTOF-Q II system) and the rates were calculated based on the buildup of cNW11 over time. The LC allowed effective separation of the unreacted fusion protein and the reaction products, and the ESI-MS confirmed the identity of each eluent and provided quantitation. In the LC step, HPLC column (Phenomenex Kinetex ® 2.6 μm C18 100 Å, 50 x 2.1 mm) was equilibrated with 80% solvent A ($H_2O$:formic acid (FA) 99.8:0.2) and 20% solvent B (ACN:H2O:FA 79.8:20:0.2). The column was maintained at 60°C in a column oven and a steady flow rate at 200 μL/min. Upon sample injection, a gradient from 20–45% solvent B was performed over 3 min, followed by a gradient from 45 to 52% solvent B over 7 min. Eluents from the column were analyzed by the ESI-MS and a series of masses corresponding to cNW11 at different m/z ratios was observed. The molecules were quantitated based on the peak area of the extracted cNW11 $[M+16H]^{16+}$ ion (*m/z* 1400.7). A calibration curve of cNW11 was generated by directly injecting 10 μL of pure cNW11 at 0.2–2 μM in reaction buffer (25 mM Tris·HCl pH 7.5 and 150 mM NaCl) onto the LC-MS. The integrated peak areas of the extracted ion were processed using Bruker Compass QuantAnalysis.

For the autocyclase system, *a*MSP11-$L_{12}$ in reaction buffer was supplemented with 1 mM BME and the reaction was initiated by the addition of 10 mM $CaCl_2$. The samples were incubated at 20°C on the LC-MS autosampler during the course of the reaction. Every 20 min, as a normalised amount, 10 μL, 5 μL or 2.5 μL of reaction mixture containing initially 2.5 μM, 5 μM or 10 μM of autocyclase, was directly injected onto the LC-MS, respectively. The reaction was monitored over 2 h and the quantity of the cNW11 product was derived from the ion peak area according to the calibration curve.

For the bimolecular system, we employed the reported linear MSP11 construct[1] which consists of an N-terminal glycine, the MSP11 sequence followed by the SrtA recognition site (LPGTG) and a C-terminal $His_6$ tag. A molar ratio of MSP:WT SrtA was maintained at 1:1 in each reaction to enable direct comparison with the autocyclase reaction. The reaction conditions were the same as described for the autocyclase reaction but at varied initial concentrations of MSP11. 10 μL, 5 μL or 2.5 μL of reaction mixture containing initially 10 μM, 20 μM or 40 μM of MSP11/SrtA were sampled on the LC-MS every 1–2 h over 10 h. All experiments were conducted in triplicates.

*Molecular dynamics*

The starting coordinates for all molecular dynamics simulations of SrtA protein with the N-terminus bound recognition sequence LPGTG were obtained from the *Staphylococcus aureus* sortase-substrate complex determined by NMR (PDB ID 2KID, model 1)[6]. The coordinates for the first two residues of the SrtA LPGTG recognition sequence were also extracted from model 1 of the 2KID NMR structural ensemble (Leu702 and Pro703). Coordinates for the remaining residues were built manually, avoiding atomic overlap and oriented towards the position of Pro703. The protonation states of the residues were determined by using PropKa at pH 7[7-9]. The three systems were energy minimised by a steepest decent algorithm in vacuum using the GROMOS 54A7[10] force field and the GROMOS11 software package[11]. Atomic position restraints (force constant 5 x $10^4$ kJ $mol^{-1}$ $nm^{-2}$) were applied to those residues obtained from the NMR SrtA structure. The initial coordinates for the manually built residues were further equilibrated with a 1 ns stochastic dynamics (SD) simulation in vacuum at 298.15 K with position restraints retained on the rest of the system. The SD simulation was performed with an atomic friction coefficient of 91 ps$^{-1}$, a 1 fs timestep, and non-bonded interactions were truncated at 1.4 nm with a reaction field correction[12] applied to electrostatic interactions beyond the cut-off.

After equilibration of the manually built residues, the systems were placed in rectangular periodic boxes such that the minimum distance to the box wall was 1.4 nm, and solvated with SPC water. Cl- ions added to neutralise the systems by replacing randomly selected water molecules. The solvated systems were equilibrated with a 10 ns molecular dynamics (MD) simulation under NPT conditions using the PMEMD simulation engine within the AMBER18 simulation package and the GROMOS 54a7[10] force field. The temperature and pressure were coupled to 295.15 K and 1 atm using Berendsen weak coupling method[13] with coupling constants of 0.1 ps and 0.5 ps respectively. The coulombic interactions were treated with the particle Mesh Ewald (PME) method and van der Waals interactions were truncated at 1.4 nm. Hydrogen containing bonds were constrained with SHAKE[14] constraint algorithm with a tolerance of $10^{-4}$ nm. Newton's equations of motion were integrated using the leapfrog algorithm with a timestep of 1 fs. The center of mass motion was removed every 5000 steps. Position restraints were retained on the Cα atoms of those residues taken from the NMR SrtA structure. A subsequent 250 ns MD simulation was performed without any position restraints applied and all other settings retained.

*Assembly of POPC nanodisc and fluorescent dye labelled nanodiscs.*

To assemble cNDs, [lipid]:[cMSP] ratios were defined based on past literature, using the equation: $N_L \times S = (0.423 \times M - 9.75)^2$, where $N_L$ is the number of lipids per ND, M is the number of amino acids in the scaffold protein and S is the mean surface area per lipid used to form the ND measured in Å$^2$ [15]. POPC was estimated to have a surface area of around 70 Å$^2$ [16]. We therefore determined that the ratios that are appropriate for assembling cNW6, cNW7, cNW9, cNW11 and cNW15 with POPC were 19:1, 30:1, 43:1, 59:1 and 282:1, respectively.

All lipids were dissolved in organic solvents, mixed if required, and evaporated to form a dry thin film before ND assembly. POPC (1-Palmitoyl-2-Oleoyl-sn-Glycero-3-Phosphocholine, Anatrace) were dissolved in chloroform, aliquoted into round bottom glass vials, evaporated using nitrogen gas to create a film and placed in a vacuum desiccator overnight. POPC and DiI (1,1'-Dioctadecyl-3,3,3',3'-Tetramethylindocarbocyanine Perchlorate, Thermo Fisher Scientific) were dissolved in chloroform separately and mixed at molar ratios of 99:1 for cNW15 assembly, and

95:5 for linear NW11, cNW11, and cNW7 assemblies. The mixed POPC-DiI solution were dried and processed as above.

To assemble POPC nanodisc, cMSP and POPC were co-dissolved at the desired ratio in reconstitution buffer (25 mM Tris·HCl pH 7.5, 100 mM NaCl, 0.5 mM EDTA and 100 mM cholate) and mixed for 1 h on ice. 0.6 g of Bio-Beads SM-2 per mL of assembly solution was added to remove detergent (cholate) and subsequently initiate ND assembly. The mixture was gently stirred for 4 h at 4 °C for complete cholate removal. The Bio-Beads were removed by filteration and the assembled discs were purified and buffer exchanged using size exclusion chromatography.  Buffer containing 20 mM Tris·HCl pH 7.5, 50 mM NaCl, 1 mM EDTA was used to equilibrate a HiLoad™ 16/600 Superdex™ 200 pg column (Cytiva) on an ÄKTA purifier FPLC and the fractions containing the cNDs were collected. The size homogeneity of cND was assessed using negative-stain transmission electron microscopy (TEM).

To assemble fluorescent dye labelled nanodiscs, The dry POPC-DiI film was dissolved in 200 µL of MSP buffer (20 mM Tris·HCl pH 7.5, 100 mM NaCl, 0.5 mM EDTA) containing 100 mM sodium cholate detergent. Freeze dried MSP protein (1 mg) was dissolved in 200 µL of MSP buffer and added to POPC-DiI solution and incubated on ice with rocking for 1 h. Biobeads (0.1 g) were added to the assembly solution every hour for 3 h, incubated on ice with rocking. The assembly solution was removed from the biobeads, filtered using a 0.45 µm syringe filter, and purified using an ÄKTA Purifier FPLC system and Superdex200 Increase 10/300 (Cytiva) size exclusion column. Fractions were collected and successful nanodisc assembly confirmed by negative stain electron microscopy. The nanodiscs were buffer exchanged into 50 mM sodium phosphate buffer pH 7.5 using an Amicon 10 kDa molecular weight cut-off centrifugal filter (Merck), and concentrated to 1 mL. The concentration of nanodiscs was determined using absorbance at 280 nm. Cyanine5-NHS ester (Lumiprobe) was dissolved in anhydrous DMSO and added to the nanodisc solution in 4X molar excess then incubated at room temperature for 5 h. Excess dye was removed by buffer exchange into phosphate buffered saline pH 7.4. The average number of DiI and Cy5 molecules incorporated into each nanodisc was calculated using absorbance (280, 546, and 649 nm), extinction coefficients of each nanodisc at 280 nm (cNW15 86582 $M^{-1}cm^{-1}$, cNW11 and linear NW11 44826 $M^{-1}cm^{-1}$, cNW7 33942 $M^{-1}cm^{-1}$), DiI at 546 nm (148000 $M^{-1}cm^{-1}$), Cy5 at 649 nm (150000 $M^{-1}cm^{-1}$), and a Cy5 correction factor at 280 nm of 0.05.

*Negative stain TEM*

The surface of carbon-coated copper grids (400-mesh) (ProSciTech Pty Ltd, Kirwan, Australia) was activated by glow-discharge for 15 s before cNDs were applied. 4 µL of cND at 20 nM (20 mM Tris·HCl pH 7.4, 100 mM NaCl and 0.5 mM EDTA) was directly pipetted onto the grid and allowed to settle for 1 min. It was followed by one wash in a drop of 1% uranyl acetate (UA) and the grid was immediately moved to a fresh drop of 1% UA to stain for an additional minute. TEM was carried out on a Hitachi HT7700 electron microscope at 80 kV.

*Expression and purification of $K_V$AP-VSD*

The plasmid encoding $K_V$AP-VSD, generated previously[17], was transformed into *E. coli* BL21-CodonPlus (DE3)-RIPL competent cells and plated on a LB agar plate with 100 µg/ml ampicillin and stored at 37°C overnight. A single colony was used to inoculate 10 mL of LB medium (containing 100 µg/ml of ampicillin) and shaken at 220 rpm and at 37°C overnight. To produce $^{15}$N-labelled $K_V$AP-VSD, the preculture was used to inoculate 2 L of LB and grown at 37°C (while shaking at 220 rpm) until the $OD_{600}$ reached ~1.0. The cells were harvested and resuspended into 1 L of modified M9 minimum media (47.74 mM $Na_2HPO_4$, 22.05 mM $KH_2PO_4$, 8.56 mM NaCl, 0.05 mM $FeCl_3$, 2 mM $MgSO_4$, 1 x MEM Vitamin Solution (Gibco), 0.4% glucose, 1 g/L $^{15}NH_4Cl$, 0.1 mM $CaCl_2$, 100 µg/ml ampicillin). The culture was grown for 0.5 h at 37°C (shaking at 220 rpm) before 0.2 mM IPTG was added to induce protein expression. The culture was further incubated for 2 h and the cells were harvested by centrifugation at 9000 x *g* for 10 min at 4 °C.

The harvested cells were resuspended in cracking buffer (50 mM Tris·HCl pH 8.0, 100 mM KCl, 20 mM DDM) at a ratio of 10 ml per gram of cell pellet. The suspension was stirred for 15 min on ice and the cells were lysed by sonication (2 x 5-min cycles of 5 s on- and 12 s off-pulse repetitions on ice; 15-min break between each cycle). The lysed cells were spun down at 30,000 x *g* for 30 min. The clear lysate was then loaded directly onto 3 mL of TALON® Superflow™ histidine-tagged protein purification resin (Cytiva) in a gravity column that was pre-equilibrated with 2 CVs of KvAP equilibration buffer (20 mM Tris·HCl pH 8.0, 100 mM KCl, 5 mM DDM). The column was washed with 4 CVs of KvAP equilibration buffer and 2 CVs of wash buffer (20 mM Tris·HCl pH 8.0, 100 mM KCl, 15 mM imidazole, 5 mM DDM) and KvAP was eluted with KvAP elution buffer (20 mM Tris·HCl pH 8.0, 100 mM KCl, 150 mM Imidazole, 5 mM DDM).

The fractions containing $^{15}N$ $K_VAP$-VSD were then purified using size exclusion chromatography using an ÄKTA purifier FPLC system. The sample was applied to a HiLoad™ 16/600 Superdex™ 200 pg column (Cytiva) that was pre-equilibrated with KvAP equilibration buffer. The flow rate was at 0.12 mL/min. The purity of the protein fractions was assessed by SDS-PAGE, the fractions containing pure KvAP-VSD were pooled and concentrated using an Amicon Ultra-15 concentrator (Merck). For cND assembly, the $^{15}N$ $K_VAP$-VSD was buffer exchanged using a PD-10 column (GE Healthcare) into 50 mM Tris·HCl pH 8.0, 100 mM KCl, 30 mM DDM.

*Nanodisc reconstitution of $^{15}N$ $K_VAP$-VSD*

To assemble $^{15}N$ $K_VAP$-VSDs into cNDs, we used purified cMSP11 in MSP buffer (20 mM Tris pH 7.5, 150 mM NaCl, 0.5 mM EDTA and 1 mM sodium azide ($NaN_3$)). 1,2-Dimyristoyl-sn-glycero-3-phosphoglycerol (DMPG; Anatrace) lipid was dissolved in chloroform:methanol:water (97:2:1), and the solvent was evaporated to form a film, and dried in a vacuum desiccator overnight. The film was solubilised in MSP buffer supplemented with 200 mM sodium cholate. The assembly mixture contained $^{15}N$-$K_VAP$-VSD, cNW11, DMPG and cholate at respective ratio of 1:20:800:1600 and the cholate concentration was maintained in the range of 12–40 mM as recommended by Ritchie et al.[15]. The materials were mixed and incubated at 25 °C with gentle rocking overnight. 0.5 mg of Bio-beads SM-2 (BIO-RAD) per mL of assembly mixture was then added , followed by shaking of the mixture at 100 rpm at 25 °C for 1.5 h for complete removal of detergents.

For the purification of $^{15}N$ $K_VAP$-VSD cNDs, the Bio-beads were removed by filtration and the cND solution was loaded onto a 1-mL Ni-NTA column (GE Healthcare) that was equilibrated with nanodisc-IMAC buffer (20 mM Tris·HCl pH 8.0, 250 mM NaCl, 1 mM sodium azide). The column was washed by 4 CVs of nanodisc-IMAC buffer and 6 CVs of nanodisc-IMAC buffer with 10 mM imidazole to remove any empty cNDs. $^{15}N$ $K_VAP$-VSD containing cNDs were eluted by 5 CVs of nanodisc-IMAC buffer with 100 mM imidazole. The eluted sample was buffer exchanged into nanodisc-NMR buffer (10 mM Tris·HCl, 10 mM EDTA, 1 mM sodium azide, pH 7.0) using a PD-10 column. The $^{15}N$ $K_VAP$-VSD cND sample was concentrated to 72 μM using an Amicon 50 kDa molecular weight cut-off centrifugal filter for subsequent NMR experiments.

*NMR experiments for KvAP and SFTI*

All NMR data were acquired using a 900-MHz NMR spectrometer equipped with a triple resonance cryogenic probe (Bruker NEO). A 2D $^{1}H$-$^{15}N$ TROSY-HSQC spectrum of 72 µM $^{15}N$-labelled $K_VAP$-VSDs in cNDs was acquired at 318 K in nanodisc-NMR buffer after addition of 5% $D_2O$.

A 200 µM of $^{13}C/^{15}N$ SFTI sample was prepared by dissolving the lyophilised peptide in 50 mM sodium acetate buffer (pH 5.2) containing 5% $D_2O$. A 2D $^{1}H$-$^{15}N$ HSQC spectrum was acquired together with 3D HNCAB, CBCAcoNH and hCCcoNH for backbone and sidechain $^{13}C$ and $^{15}N$ resonance assignments – all at 298 K. All 3D experiments were acquired using non-uniform sampling[18] and processed using automated maximum entropy reconstruction[19].

*Biodistribution of fluorescent nanodiscs*

This study was carried out in accordance with the recommendations of Animal Care and Protection Act (2001) and the Australian code for the care and use of animals for scientific purposes (8th Edition). The protocol was approved by a University of Queensland Anatomical Bioscience Ethics Committee (Approval AIBN/105/19). Imaging experiments were performed using a IVIS Lumina X5 imaging system (Perkin Elmer) to visualise DiI (540 nm excitation, 620 emission filters) and Cy5 (620 nm excitation, 670 emission filters). Naïve female BALB/c nude mice were anesthetised with 3–5% isoflurane and injected intravenously via the tail vein with 150 µL of nanodisc solution (3 µM in phosphate buffered saline pH 7.4). The animals were anesthetised during image acquisition and allowed to recover after each imaging time point. The mice were sacrificed 24 hours post injection, and the major organs were removed for ex vivo imaging. Images were analyzed using Living Image Software (Perkin Elmer). For ex vivo images, regions of interest (ROI) were defined around each tissue. The average radiant efficiency ([p/s/cm$^2$/sr]/[µW/cm$^2$]) of each ROI minus the average radiant efficiency of the relevant non-injected control tissue was determined and normalised to blood. Unpaired t-test statistical analysis was performed using GraphPad Prism software (version 8.3.1).

#### Supplementary Figures

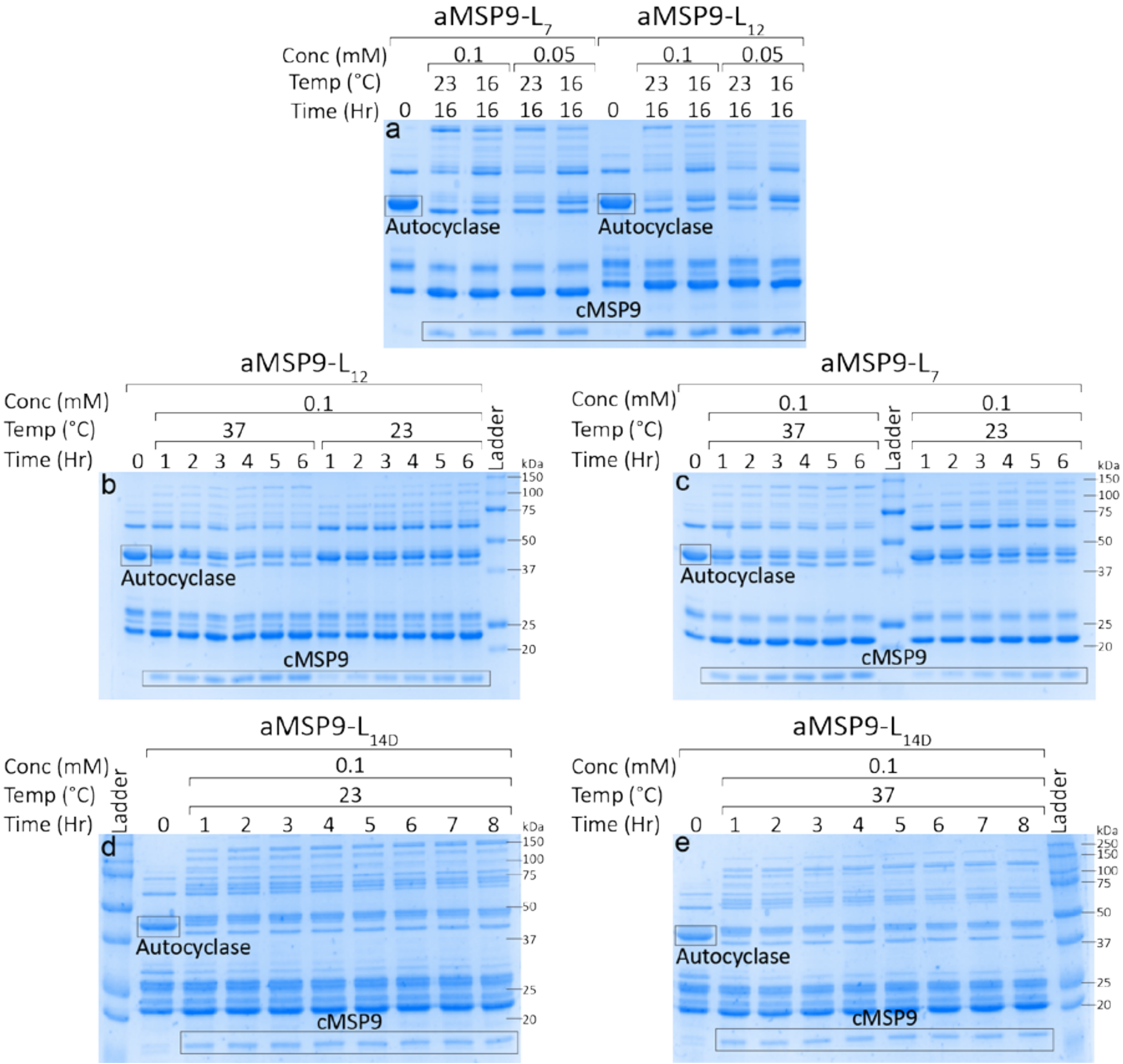


**Supplementary Figure 1: Catalytic efficiency of autocyclase cyclisation is dependent on the linker that connects the SrtA recognition site to the N-terminus of SrtA.** (Data used in Fig. 1c and 1d). The cyclisation reactions of *a*MSP9 with the L$_7$, L$_{12}$ and L$_{14D}$ linkers were monitored using SDS-PAGE. Reaction variables include initial autocyclase concentration (0.05 or 0.1 mM), temperature (37°C or 23°C) and reaction time (1-16 h). All reactions here were supplemented with 1 mM DDM. The amount of sample loaded per lane was normalised according to the initial protein concentration. Lanes denoted with "Ladder" were loaded with molecular weight standard. Bands corresponding to the unreacted autocyclase and cMSP9 products are labelled and highlighted by black boxes. **a,** *a*MSP9-L$_7$ and *a*MSP9-L$_{12}$ reactions carried out at 0.1 or 0.05 mM at 23°C or 16°C for 16 hours. **b,** 0.1 mM *a*MSP9-L$_{12}$ reaction at 37°C or 23°C over 6 hours. **c,** 0.1 mM *a*MSP9-L$_7$ reaction at 37°C or 23°C over 6 hours. **d,** 0.1 mM *a*MSP9-L$_{14D}$ reaction at 23°C over 8 hours. **e,** 0.1 mM *a*MSP9-L$_{14D}$ reaction at 37°C over 8 hours.

We perform the cyclisation reaction using fusion proteins directly after Ni-NTA purification at the concentration of 0.05 mM or 0.1 mM. At higher concentrations, MSPs tend to oligomerise. Thus, we can conclude that the extra bands are either due to MSP9 oligomers or impurities carried from Ni-NTA purification (*E. coli* proteins and/or fusion-protein truncations). However, we focus on measuring the intensities of the gel bands corresponding to cMSP9 and *a*MSP, highlighted by the rectangles – measure is independent of the presence of impurities.

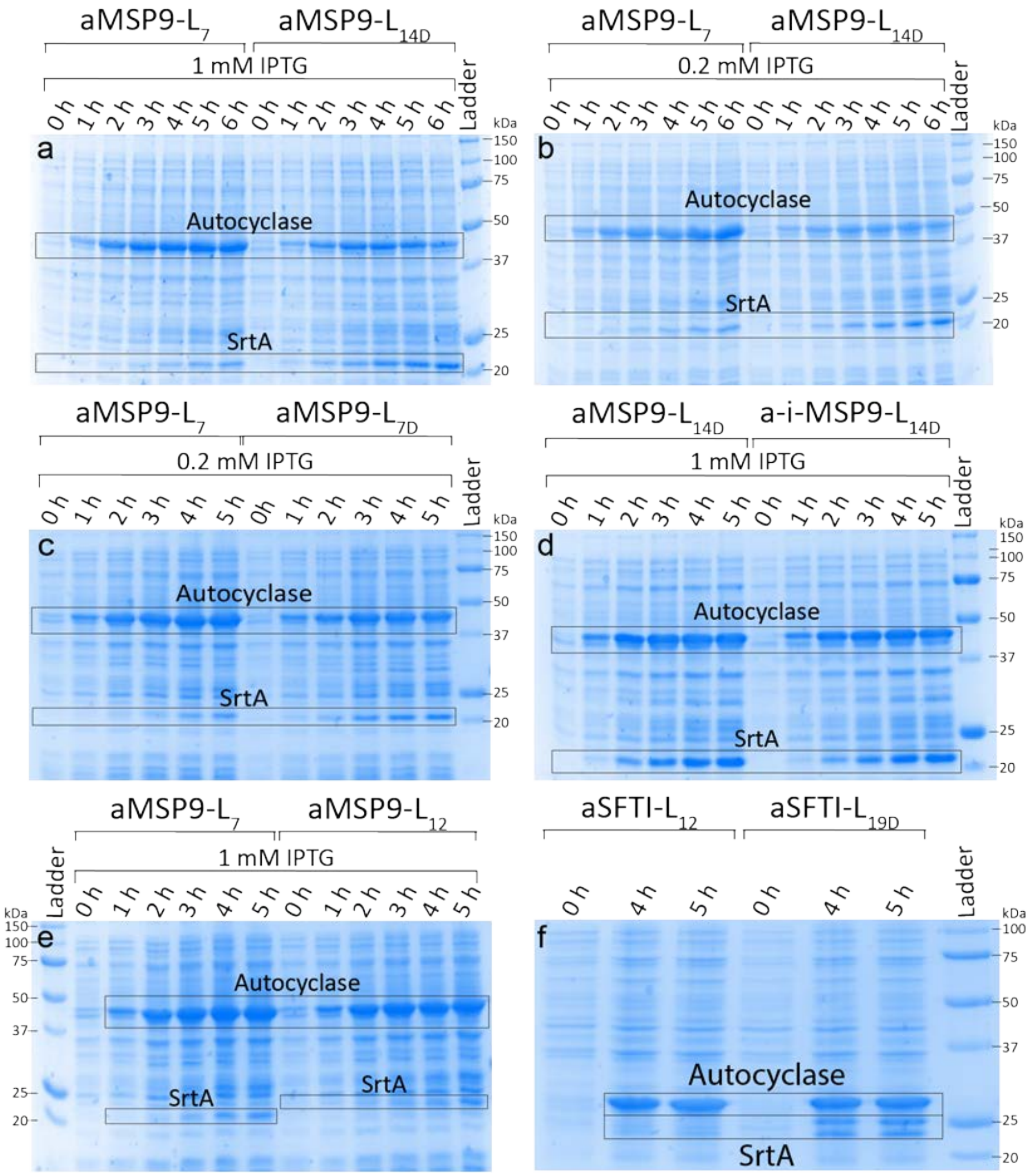


**Supplementary Figure 2: Linkers that promote disorder are prone to in vivo hydrolysis during recombinant bacterial expression.** Expression of autocyclases in *E. coli* was monitored using SDS-PAGE. Cells were sampled every hour post-induction over 5-6 hours and in vivo hydrolysis of the autocyclase linker is evident by the gradual build-up of SrtA during the course of expression. Long and more dynamic linkers appeared to be more susceptible to this in vivo degradation as judged by the more intense SrtA band on the gel (relative to the autocyclase band). **a,** Lysed cell suspension sampled hourly to compare expression of *a*MSP9-$L_7$ and -$L_{14D}$ (induced with 1 mM IPTG). **b,** The same as **a** but induced using 0.2 mM IPTG. **c,** Same as **b** but comparing *a*MSP9-$L_7$ and *a*MSP9-$L_{7D}$. **d,** Same as **a** but comparing *a*MSP9-$L_{14D}$ with a variant that contains an N-terminal inhibitory peptide (*a*-*i*-MSP9-$L_{14D}$). **e,** Same as **a** but comparing *a*MSP9-$L_7$ and *a*MSP9-$L_{12}$. **f,** In vivo hydrolysis was also observable (but weaker) when using disorder promoting linkers for non-MSP autocyclases — here comparing *a*SFTI-$L_{12}$ and *a*SFTI-$L_{19D}$.

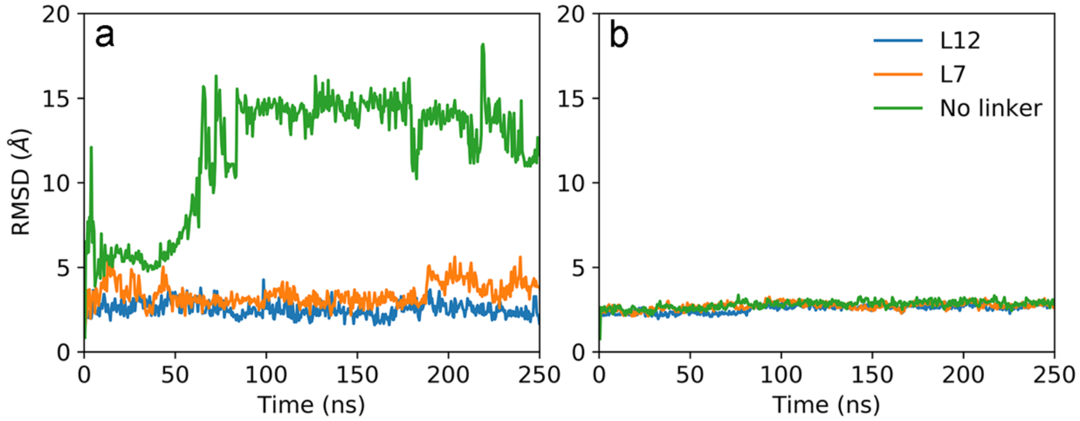


**Supplementary Figure 3: The positional root mean squared deviation (RMSD) from molecular dynamics simulations.** The simulations were performed using a sortase-substrate complex with the recognition sequence attached to the N-terminus of SrtA with the $L_{12}$ linker (blue), $L_7$- linker (orange) and no-linker (green). **a**, shows the RMSD of the LP residues within the recognition sequence (LPGTG). **b**, shows RMSD of all residues involved in secondary structure (PDB ID: 2KID). In all cases RMSD fits were performed on secondary structure residues with respect to the starting configuration derived from model 1 of PDB ID: 2KID.

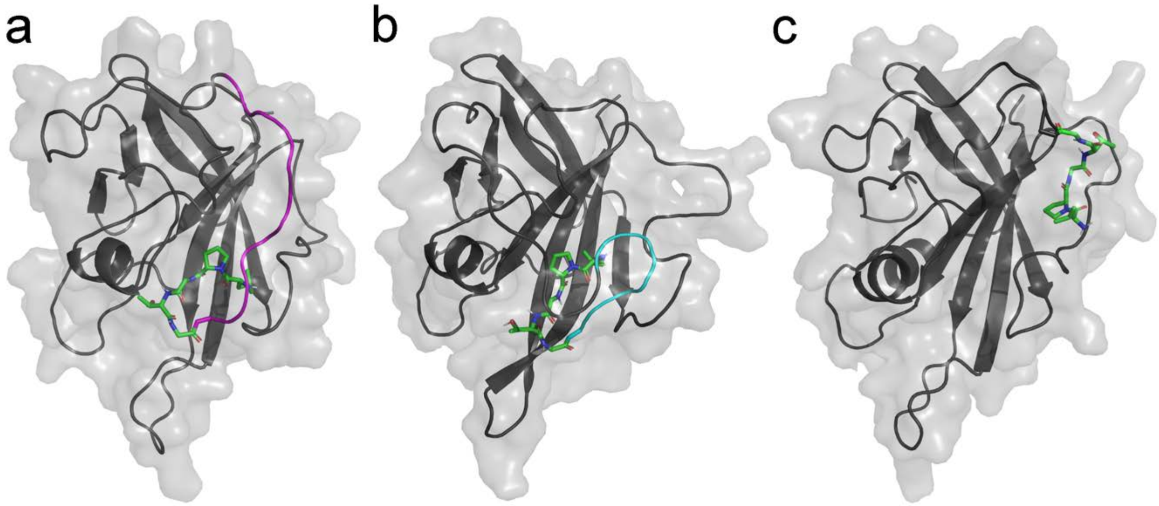


**Supplementary Figure 4: Snapshots of the final structure from 250 ns molecular dynamics simulations of Sortase A.** SrtA structure (cartoon) with substrate (sticks) attached to the N-terminus with: $L_{12}$- linker (magenta, Panel **a**), $L_{7}$- linker (cyan, Panel **b**) and no-linker (Panel **c**).

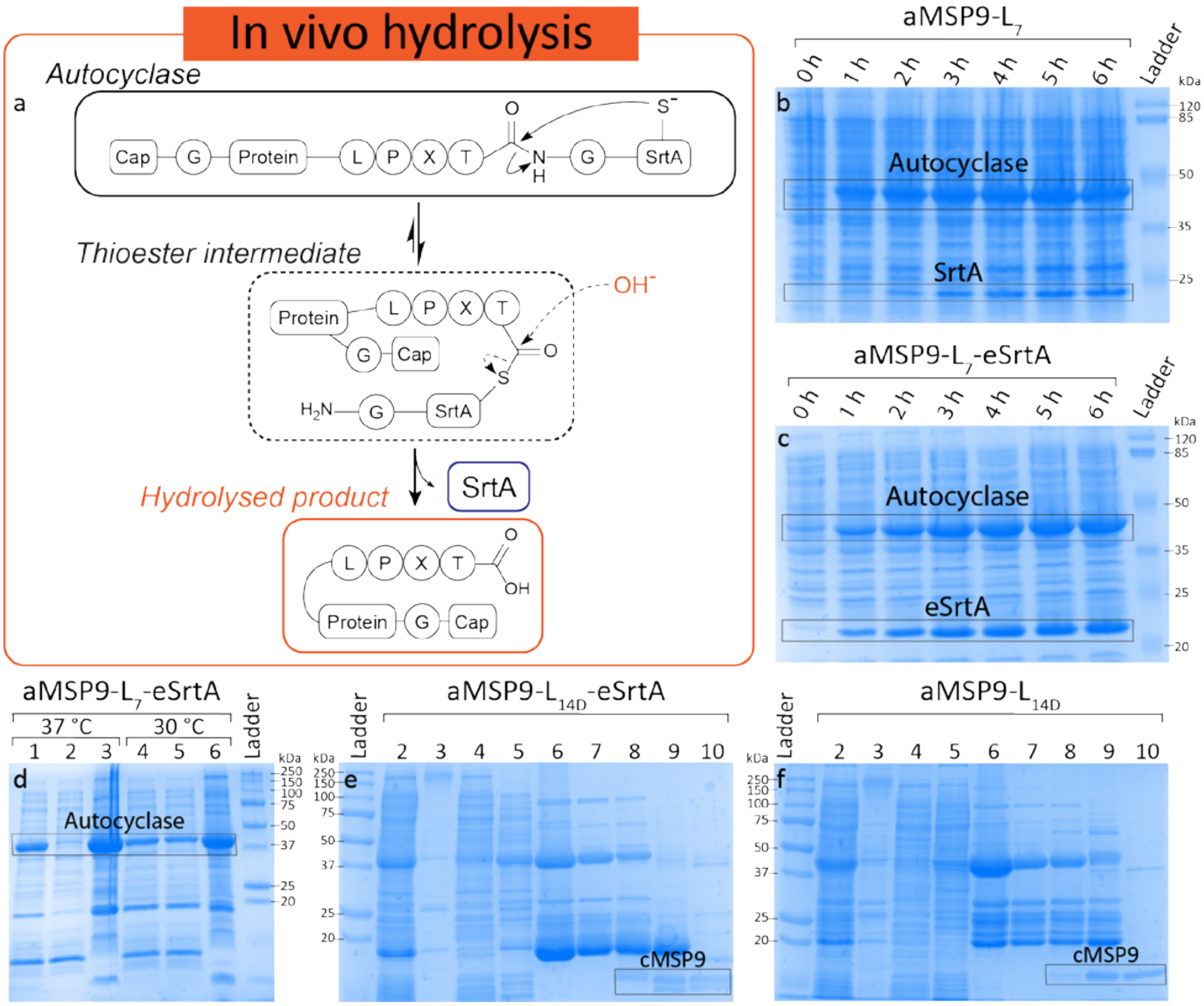


**Supplementary Figure 5: Autocyclases containing the evolved sortase (pentamutant - eSrtA) are less soluble and prone to in vivo hydrolysis resulting in poor overall yields.**
**a,** A schematic illustrating the pathway of autocyclase hydrolysis in vivo. **b-c,** Comparison of the expression of *a*MSP9-$L_7$ (-WT SrtA; b) and *a*MSP9-$L_7$-eSrtA (c) in *E. coli* monitored using SDS-PAGE. Cells were sampled every hour post-induction over 6 hours and in vivo hydrolysis of the autocyclase was monitored by observing the gradual build-up of SrtA/eSrtA during the course of expression. **d,** Expression of *a*MSP9-$L_7$-eSrtA at 37 (lane 1-3) or 30°C (lane 4-6). The soluble (lane 2 and 5) and insoluble (lane 3 and 6) fractions extracted from cells expressing *a*MSP9-$L_7$-eSrtA (whole cells in lane 1 and 4) show that the eSrtA-fused autocyclase was insoluble when it was expressed at higher temperature—suggesting poor thermal stability. **e-f,** The process of producing cMSP9 from *a*MSP9-$L_{14D}$-eSrtA (e) and *a*MSP9-$L_{14D}$-SrtA (f) was monitored by SDS-PAGE. Samples were collected at each step of the process: (i) cell lysis—soluble (lane 2) and insoluble (lane 3) fractions, (ii) Ni-NTA purification—column flow-through (lane 4), low imidazole concentration wash (lane 5) and elution (lane 6), (iii) cyclisation reaction—before reaction (lane 7), 1 h after reaction (lane 8), 15 h after reaction (lane 9) and post-reaction purification using reverse-IMAC (lane 10). It is apparent that autocyclases containing eSrtA are more prone to yielding side products during the autocyclisation reaction.

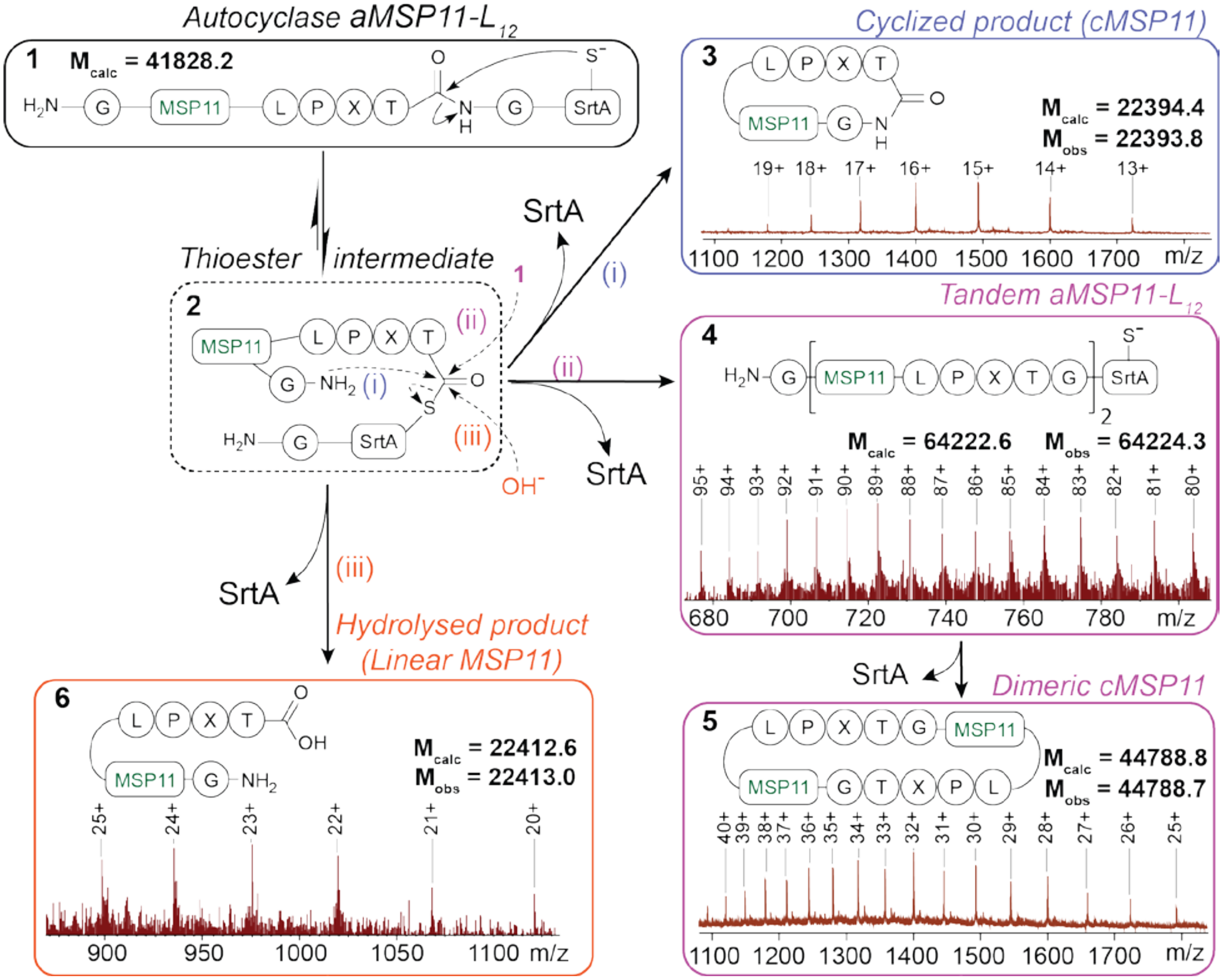


**Supplementary Figure 6: Support of the proposed reaction pathways of autocyclases by mass spectrometry. a,** A schematic illustrating the three possible reaction fates of autocyclases. Each reaction was supported by the identification of masses that correspond to intermediates and/or products by mass spectrometry. The reaction was carried out using 50 µM *a*MSP11-L$_{12}$ at room temperature overnight. Pathway (i) produces the desired cMSP11 as confirmed by a series of observed *m/z* ratios including 1318.3 $[M+17H]^{17+}$ and 1400.6 $[M+16H]^{16+}$. Pathway (ii) produces a tandem *a*MSP11-L$_{12}$ that yields a dimeric cMSP11. Masses (*m/z*) consistent with both molecules could be identified from the reaction mixture. They include 714.6 $[M+90H]^{90+}$ and 722.6 $[M+89H]^{89+}$ for the tandem autocyclase and 1318.3 $[M+34H]^{34+}$ and 1358.2 $[M+33H]^{33+}$ for the dimer. Pathway (iii) produced linear MSP11 via hydrolysis. *m/z* consistent with linear MSP11 including 975.5 $[M+23H]^{23+}$ and 1019.8 $[M+22H]^{22+}$ were identified.

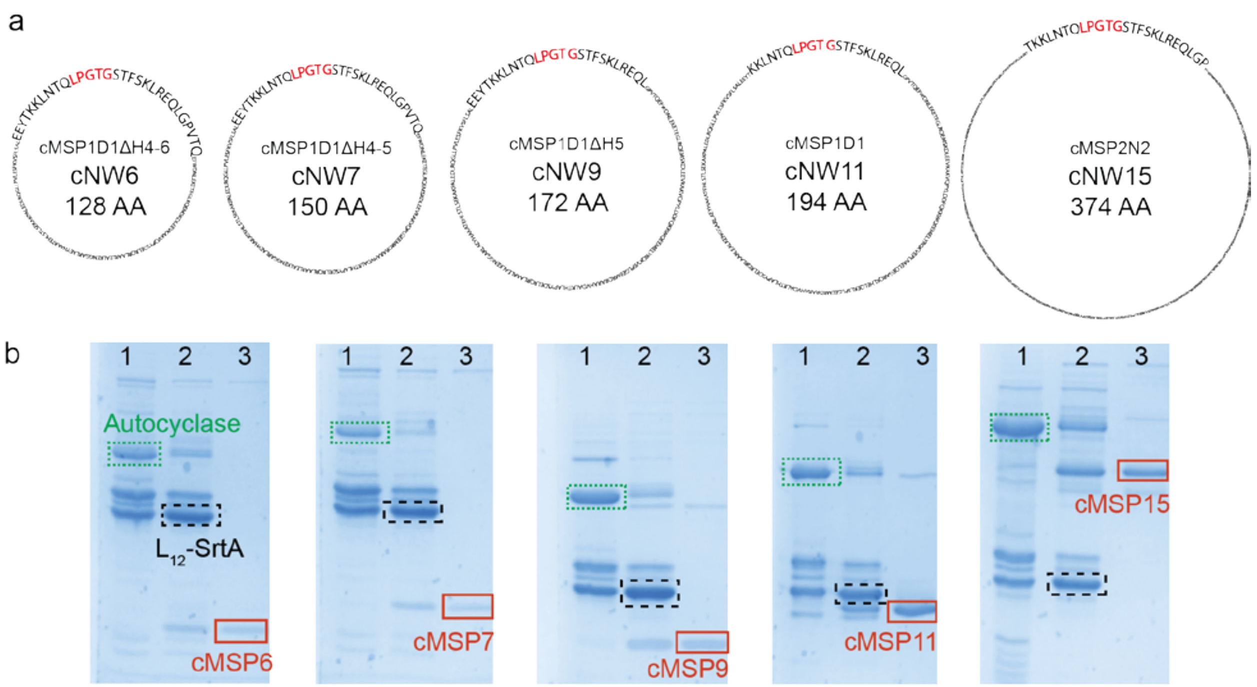


**Supplementary Figure 7: The production of different sizes of cyclic membrane scaffold proteins (cMSPs) from autocyclases. a,** the diagrams of cMSP6, 7, 9, 11 and 15 (number indicates diameter of assembled nanodisc in nm). **b,** SDS-PAGE images illustrating the production of cMSPs (concentration ~50-100 µM, reaction at 37°C, ~18 h). Lane 1 and 2 of each gel shows the reaction mixtures at the beginning and the end of the reaction, respectively. Lane 3 shows the final cMSP products after reverse-IMAC purification. The unreacted autocyclases are highlighted in green boxes, SrtA produced from the reaction (still attached to the linker) in black boxes and the cMSP products in red boxes. To confirm cyclisation, the exact molecular weight of each cMSP was validated by ESI-MS (Supplementary Fig. 7). A distinctive 18 Da difference can be observed between the linear and cMSP. The cMSP6 and cMSP7 bands move relatively slowly in the 15% SDS-PAGE gel compared to the others (cMSP9, 11 and 15) that were run using a 12% gel. The autocyclase used in the reaction were performed directly after Ni-NTA purification and the samples were at ~60% purity. It is apparent that the reaction does not require the sample to be of high purity to proceed and the product will subsequently be purified post-reaction.

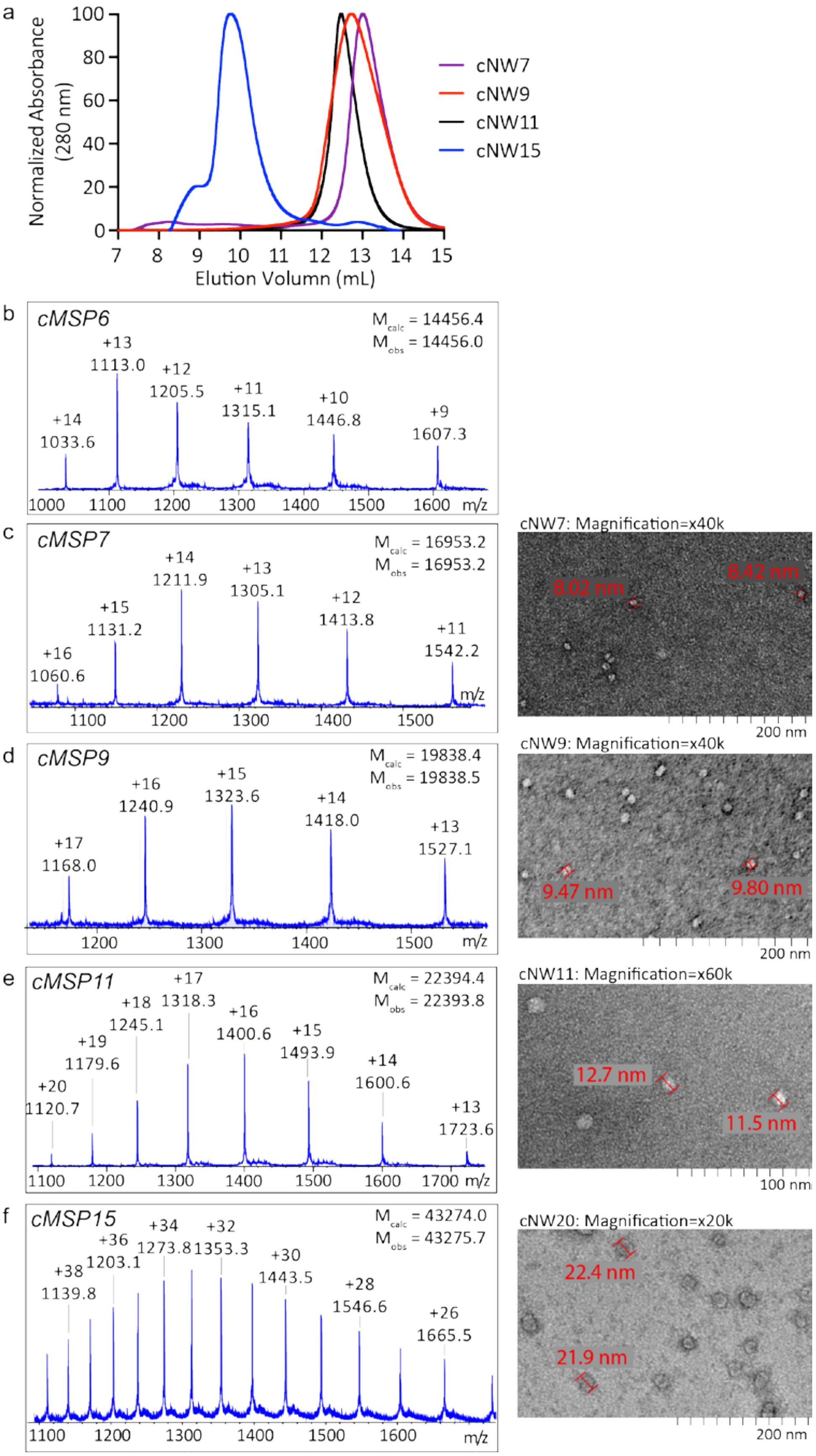

a
Normalized Absorbance (280 nm)
Elution Volumn (mL)
cNW7
cNW9
cNW11
cNW15
b
cMSP6
M calc = 14456.4
M obs = 14456.0
+14 1033.6
+13 1113.0
+12 1205.5
+11 1315.1
+10 1446.8
+9 1607.3
m/z
c
cMSP7
M calc = 16953.2
M obs = 16953.2
+16 1060.6
+15 1131.2
+14 1211.9
+13 1305.1
+12 1413.8
+11 1542.2
m/z
d
cMSP9
M calc = 19838.4
M obs = 19838.5
+17 1168.0
+16 1240.9
+15 1323.6
+14 1418.0
+13 1527.1
m/z
e
cMSP11
M calc = 22394.4
M obs = 22393.8
+20 1120.7
+19 1179.6
+18 1245.1
+17 1318.3
+16 1400.6
+15 1493.9
+14 1600.6
+13 1723.6
m/z
f
cMSP15
M calc = 43274.0
M obs = 43275.7
+38 1139.8
+36 1203.1
+34 1273.8
+32 1353.3
+30 1443.5
+28 1546.6
+26 1665.5
m/z
cNW7: Magnification=x40k
8.02 nm
8.42 nm
200 nm
cNW9: Magnification=x40k
9.47 nm
9.80 nm
200 nm
cNW11: Magnification=x60k
12.7 nm
11.5 nm
100 nm
cNW20: Magnification=x20k
22.4 nm
21.9 nm
200 nm

**Supplementary Figure 8: Confirmation of cMSP6, 7, 9, 11 and 15 production by mass spectrometer and assembly of cyclised POPC-nanodiscs cNW7, 9, 11 and 15. a,** Size exclusion chromtograms of POPC-cNW7 (purple), 9 (red), 11 (black) and 15 (blue) showing the different elution time of the cNDs. **b-f,** The mass of cMSP 6 (b), 7 (c), 9 (d), 11 (e) and 15 (f) were confirmed using mass spectrometry. All observed masses (*m/s)* are consistent with the calculated masses of the cyclised products. The cMSP7-15 were then used to generate nanodiscs that encircled POPC lipid bilayers. Negative stain TEM images of the cNDs are shown on the right of the corresponding cMSP mass spectra. Estimated diameter of the disc are included to show that the cNDs adopt the expected size.

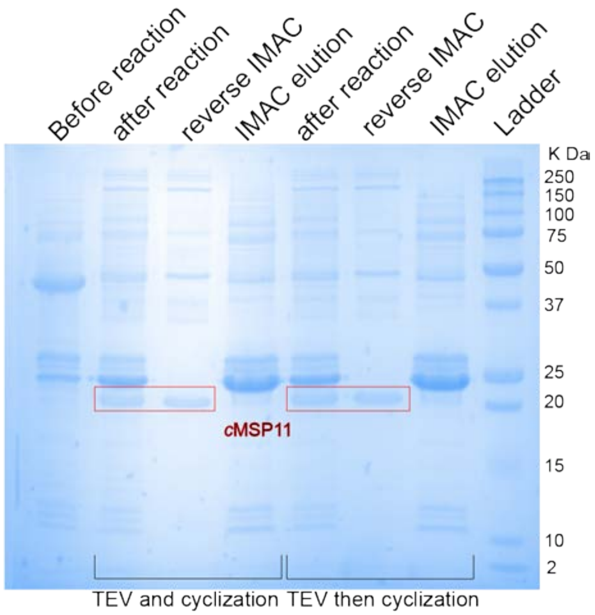


**Supplementary Figure 9. cMSP11 production by TEV protease cleavage and cyclisation at the same time.** After IMAC purification and buffer exchange, *a*MSP11 (~50 kDa) is subjected to TEV protease cleavage and cyclisation simultaneously overnight, which readily produces cMSP11 (red box). cMSP11 production is identical between the process where *a*MSP11 is subjected to TEV protease cleavage and cyclisation together and the process where *a*MSP11 undergoes TEV protease cleavage first, followed by cyclisation.

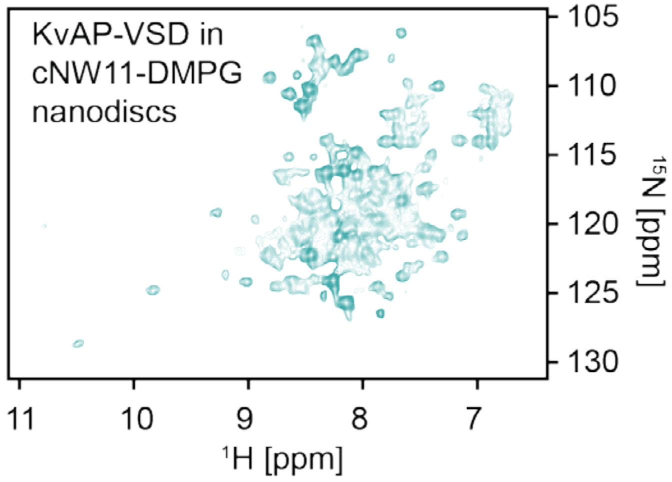


**Supplementary Figure 10: $^{1}H$-$^{15}N$ TROSY spectrum of $^{15}N$-labelled KvAP-VSD in POPC-cNW11.** NMR spectrum of the correctly folded protein in detergent micelles and non-cyclised ND have been reported[20], and agree well with the spectrum measured here.

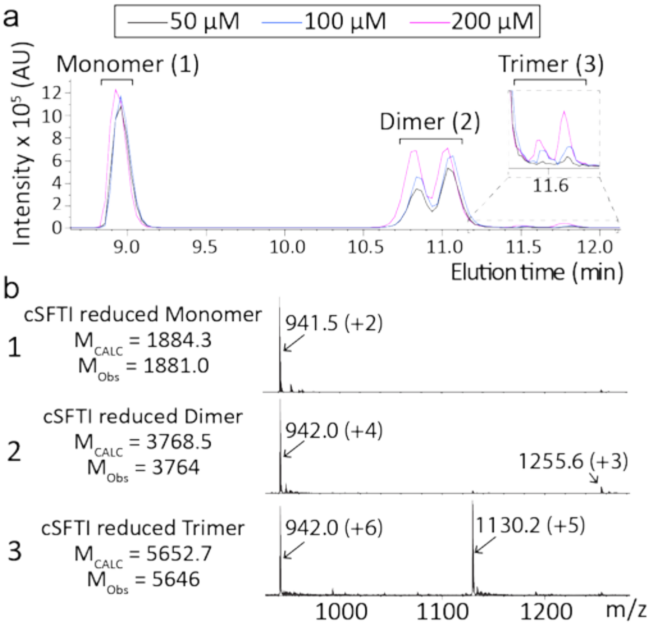


**Supplementary Figure 11: Polymerisation of SFTI autocyclisation.** Cyclisation of *a*SFTI-L$_{14D}$ at initial concentrations of 50 (black), 100 (blue) and 200 µM (pink) was assessed using LC/MS. The reaction was carried out at 37°C overnight. **a,** the chromatogram of the extracted ions with a *m/z* of 941 is illustrated. A *m/z* of 941 corresponds to the $[M+2H]^{+2}$, $[M+4H]^{+4}$ or $[M+6H]^{+6}$ ion of monomeric, dimeric or trimeric cSFTI, respectively. **b,** *m/z* ions identified from the peaks at 9, 11 and 11.8 min confirmed the oligomeric states of the cSFTI to be monomeric, dimeric and trimeric, respectively. Reactions carried out at higher initial concentrations lead to a higher proportion of polymers in a concentration dependent manner.

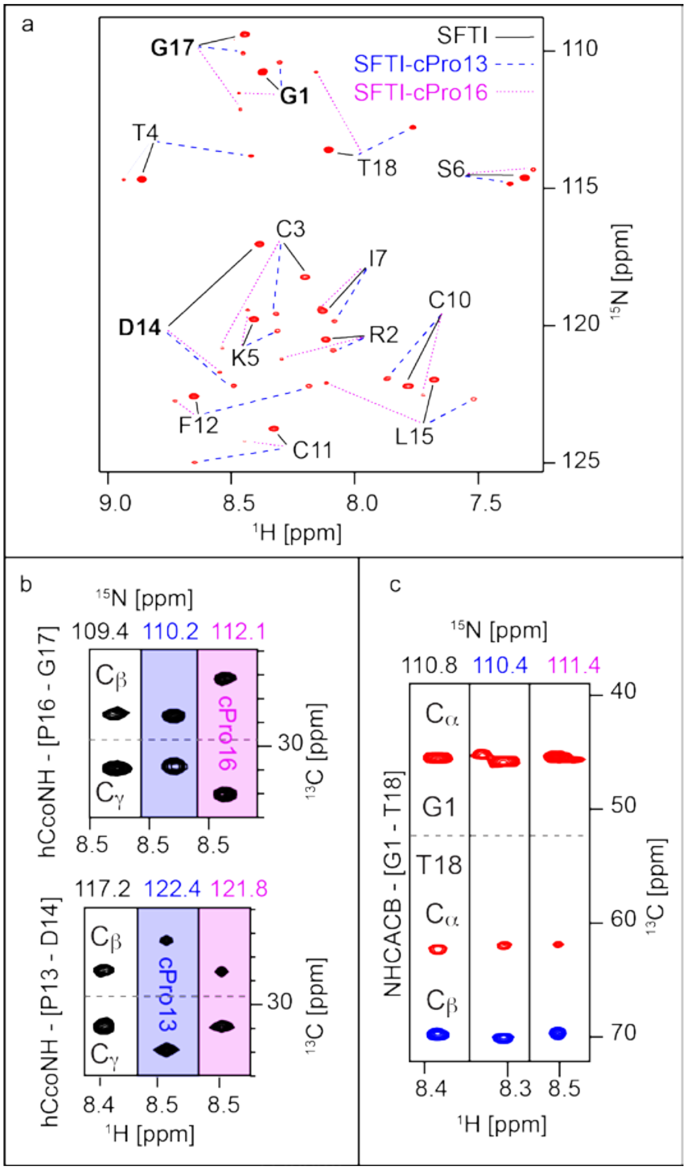


**Supplementary Figure 12: NMR data of autocyclised $^{13}C/^{15}N$ (LPGTG)SFTI. a**, Assigned $^{1}H$-$^{15}N$ HSQC spectrum showing that SrtA cyclised SFTI exists in 3 stable conformations in solution. **b**, Strip-plots taken at the $^{1}H$-$^{15}N$ chemical shift of the residue following proline residues (P13 and P16), D14 (bottom) and G17 (top) respectively, where correlations to the proline $^{13}C$ atoms can be observed. Each set of three strip-plots is taken at the chemical shift corresponding to the three different conformations. The three conformations arise from cis/trans isomerisation of Pro-13 and Pro-16 as indicated. The difference in $^{13}C$ chemical shift between the $C_\beta$ and $C_\gamma$ is distinctly larger where the proline is in a cis conformation (cPro). **c**, In the HNCACB experiment there is a clear signal from the HN resonances of Gly-1 to the $C_\alpha$ and $C_\beta$ atoms of Thr-18. These correlations are due to scalar couplings between covalently bound atoms, providing clear evidence of head-to-tail macrocyclisation in each of the three conformations of SFTI in solution.

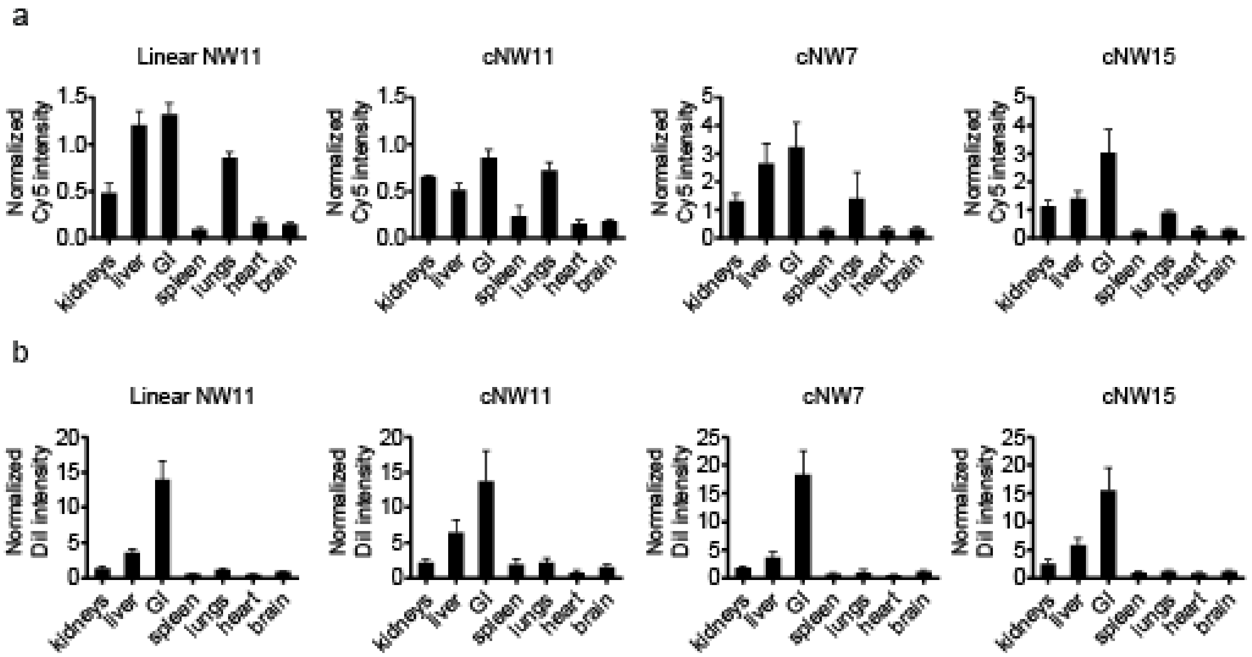


**Supplementary Figure 13: Biodistribution of nanodiscs in naïve BALB/c nude mice.** Animals administered fluorescent nanodiscs were euthanised 24 hours post injection. Organs were immediately dissected and imaged using 540 nm excitation/620 nm emission filters (DiI) and 620 nm excitation/670 nm emission filters (Cy5). **a,** biodistribution of Cy5-conjugated MSP proteins. **b,** biodistribution of DiI lipid dye. Fluorescence data is presented as mean normalised (to blood) radiant efficiency values (n=3) with standard deviation error bars.

### Supplementary Tables

**Supplementary Table 1: The average yield of purified autocyclase per liter of bacterial culture.** All yields are averages of triplicates of 300 mL bacterial cultures grown in LB media. The data is then normalised to 1-liter scale. The yields are calculated by the following process. After Ni-NTA purification and buffer exchange, we measure the absorbance of the purified proteins at 280 nm to calculate the overall mass of the eluted proteins (using the autocyclase extinction coefficient). We then ran the purified proteins on an SDS-page gel and measured the band intensity of autocyclase relative to all bands to estimate the abundance of autocyclase (relative to other bands). The major contaminant was found to be free SrtA as a result of in vivo processing.

| Autocyclase | Linker | % Fusion (subtracting free SrtA) | Autocyclase MW | cMSP/cyclic peptide MW | Average yield (mg) | Average yield (mmol) |
|---|---|---|---|---|---|---|
| MSP6 | $L_{12}$ | 24±1% | 34930.3 | 14456.4 | 18.6±1.4 | 0.5±0.0 |
| MSP7 | $L_{12}$ | 16±3% | 37427.2 | 16953.2 | 12.4±4.0 | 0.3±0.1 |
| MSP9 | $L_{12}$ | 39±2% | 40312.4 | 19838.4 | 63.6±3.4 | 1.6±0.1 |
| MSP11 | $L_{12}$ | 46±3% | 42868.3 | 22394.4 | 65.0±14.4 | 1.5±0.3 |
| MSP15 | $L_{12}$ | 55±1% | 63747.9 | 43274.0 | 60.0±11.8 | 0.9±0.2 |
| SFTI | $L_{19D}$ | 45±1% | 23246.1 | 1884.3 | 109.2±7.1 | 4.7±0.3 |
| kB1 | $L_{19D}$ | 45±1% | 24418.3 | 3056.5 | 83.1±4.6 | 3.4±0.2 |
| Vc1.1 | $L_{19D}$ | 49±2% | 23085.7 | 2276.5 | 118.1±3.6 | 5.1±0.2 |

**Supplementary Table 2: Cyclisation efficiency of different sizes of MSPs produced by the corresponding autocyclase.** Reactions performed at 5 µM starting concentration of pure autocyclase (following anion exchange) at 37°C for 18 h (data are averages of 3 reactions).

| **Autocyclase** | **Cyclisation reaction (Monomeric cMSP recovery)** | **Estimated yield of cMSP per litre of expression*** | | |
|---|---|---|---|---|
| | Average cyclisation yield (%) | Yield (mg) | Yield (mmol) | Yield (mg) from reported bimolecular methods[1, 21] ** |
| *a*MSP6-$L_{12}$ | 89 ± 8 | 7 | 0.5 | N/A |
| *a*MSP7-$L_{12}$ | 90 ± 5 | 5 | 0.3 | N/A |
| *a*MSP9-$L_{12}$ | 90 ± 3 | 28 | 1.4 | 12 |
| *a*MSP11-$L_{12}$ | 81 ± 5 | 28 | 1.2 | 14 |
| *a*MSP15-$L_{12}$ | 90 ± 4 | 33 | 0.8 | 16 |

**Estimated based on yield of pure autocyclase (per liter of expression – from Table S1) and the observed cyclisation efficiency.*

***Cyclisation yields by bimolecular method have only been reported by Yusuf et al.[21].*

**Supplementary Table 3: Primers used in this study.**

| Primer number and descriptions | Primer sequences |
|---|---|
| P1-Forward with *Kpn*I sites underlined, for SrtA | CGG GAATCC GGTACC CAA GCTAAACCTCAAATT CCGAAAG |
| P1-Reverse with *Xho*I site underlined | TTTTTT CCG CTCGAG TTT GAC TTC TGT AGC TAC AAA GAT TTT ACG |
| P2-forward, for removing N-terminal His-tag | GAGAATTTGTACTTCCAAGGATC |
| P2-Reverse | GGACGAAGCCATATGTATATC |
| P3-forward, for introducing His4 into His$_6$ at the C-terminal | CATCATTGAGATCCGGCTGCTAAC |
| P3-reverse | ATGATGGTGGTGGTGGTGGTGGTG |
| P4-forward, for introducing GS(GGS)$_4$ linker ($L_{14D}$) in MSP9-eSrtA | TGGGTCCGGTGGTAGTGGTGGGAGTCAAGCTAAACCGCAGATC |
| P4-reverse | CCTGAACCTCCCGATCCCCCGGTACCAGGCAGTTGTGTGTTAAG |
| P5-forward, for introducing GS(GGS)$_4$ linker ($L_{14D}$) in MSP9-SrtA | TGGGTCCGGTGGTAGTGGTGGGAGTCAAGCTAAACCTCAAATTC |
| P5-reverse = P5-reverse | CCTGAACCTCCCGATCCCCCGGTACCAGGCAGTTGTGTGTTAAG |
| P6-forward with *Nde*I site underlined, for MSP11 | CGCGGATCC CAT ATG GCT AGC AGC GAA AAC CTG TAT TTT CAG GGC AGC ACC |
| P6-reverse with *Kpn*I site underlined | GGCGAATTC GGT ACC CGG CAG CTG GGT G |
| P7-forward, for deleting H4 in MSP9 to make MSP7 | TTGGGGGAGGAGATGCGT |
| P7-reverse | GGGTTGTACCTTAGCCTTCAC |

| | |
|---|---|
| P8-forward, for deleting H4 and H6 in MSP9 to make MSP6 | TATAGTGATGAGTTGCGC |
| P8-reverse | GGGTTGTACCTTAGCCTTC |
| P9-forward, optimised inhibitory peptide | CGCAGAGAATTTGTATTTCCAGGGATCGACGTT TTCCAAG |
| P9-reverse | TCGCGTGGAAGGGAGGAAGCCATATGTATATCT CCTTCTTAAAGTTAAAC |
| P10-forward, for introducing a Thrombin site (LVPRS) between LPGTGAAALEGT linker and SrtA | GCGCAGCCAAGCTAAACCTCAAATTCC |
| P10-reverse | GGCACCAGGGTACCCTCTAAAGCTGC |
| P11-forward, for introducing a Thrombin site (LVPRS) between LPGTG(GGS)$_5$ linker and SrtA | GCGCTCCCAAGCTAAACCTCAAATTCCGAAAG |
| P11-reverse | GGCACGAGACTCCCACCACTACCACC |
| P12-forward, for removing N-terminal his tag in MSP11-LPGTGAAALEGTLVPRS-SrtA-His$_{10}$ | GAAAACCTGTATTTTCAGGG |
| P12-reverse | GCTGCTAGCCATATGTATATC |
| P13-forward, for amplification of MSP15 and replace MSP9 in MSP9-LPGTGAAALEGTLVPRS-SrtA-His10 | AAGAAGGAGATATACATATG GCCAGTTCT<br>GAAAACCTGTATTTTCAGGGATCGACG |
| P13-reverse | CAC CAG GGT ACC CTC TAA AGC TGC AGC ACC TGT ACC AGG TAA CTG TGT ATT TAA CTT TTT AGT ATA TTC TTC |
| P14-forward, for generating empty autocyclase-L$_{19D}$ vector | CTGCCTGGTACCGGGGGA |
| P14-reverse | TCCCTGAAAATACAGGTTTTCCGCG |
| P15-forward, to generate autocyclase-L$_{19D}$-G-SFTI | GCCGATTTGCTTTCCGGATCTGCCTGGTACCGG GGGA |

| | |
|---|---|
| P15-reverse | GGAATGCTTTTGGTGCAGCGTCCCTGAAAATACAGGTTTTCCGCG |
| P16-forward, to generate autocyclase-$L_{19D}$-G-KalataB1 | CACCTGCAGCTGGCCGGTGTGCACCCGCAACGGCCTGCCGGTGACCGGGGGATCGGGAGGT |
| P16-reverse | CAGCCCGGGGTGTTGCAGGTGCCGCCCACGCAGGTTTCGCCGCATCCCTGAAAATACAGGTTTTCCGCG |
| P17-forward, to generate autocyclase-$L_{12}$-G-SFTI | GCCGATTTGCTTTCCGGATCTGCCTGGCACAGGTGCT |
| P17-reverse | GGAATGCTTTTGGTGCAGCGTCCTTGGAAGTACAAATTCTCGGAC |
| P18-forward, to generate autocyclase-$L_{12}$-GG-Vc1.1 | CTATGATCATCCGGAAATTTGCGGTCTGCCTGGCACAGGTGCT |
| P18-reverse | TTGCAGCGCGGATCGCTGCAGCAACCTCCTTGGAAGTACAAATTCTCGGAC |
| P19-forward, to generate autocyclase-$L_{19D}$-GG-Vc1.1 | TGGGTCCGGTGGTAGTGGTGGGAGTCTGGTGCCGCGCAGCCAA |
| P19-reverse | CCTGAACCTCCCGATCCCCCGGTACCAGGCAGACCGCAAATTTCCGG |

**Supplementary Table 4. Autocyclase constructs produced in this study.** The corresponding sequences are provided in Supplementary Data 1 where restriction enzyme sites are highlighted in yellow and TEV cleavage sites, SrtA recognition sequences and linkers are highlighted in grey.

| **Construct no.** | ***Name*** | **Description** |
|---|---|---|
| 1 | *a*MSP6-$L_{12}$ | MSP6-LPGTG-$L_{12}$-SrtA-$His_{10}$ |
| 2 | *a*MSP6-$L_{14D}$ | MSP6-LPGTG-$L_{14D}$-SrtA-$His_{10}$ |
| 3 | *a*MSP7-$L_{12}$ | MSP7-LPGTG-$L_{12}$-SrtA-$His_{10}$ |
| 4 | *a*MSP7-$L_{14D}$ | MSP7-LPGTG-$L_{14D}$-SrtA-$His_{10}$ |
| 5 | *a*MSP9-$L_7$-eSrtA | MSP9-LPGTG-$L_7$-eSrtA-$His_{10}$ |
| 6 | *a*MSP9-$L_7$ | MSP9-LPGTG-$L_7$-SrtA-$His_{10}$ |
| 7 | *a*MSP9-$L_{7D}$ | MSP9-LPGTG-$L_7$D-SrtA-$His_{10}$ |
| 8 | *a*MSP9-$L_{12}$ | MSP9-LPGTG-$L_{12}$-SrtA-$His_{10}$ |
| 9 | *a*MSP9-$L_{14D}$ | MSP9-LPGTG-$L_{14D}$-SrtA-$His_{10}$ |
| 10 | *a*-*i*-MSP9-$L_{14D}$ | Inhibitory Sequence-MSP9-LPGTG-$L_{14D}$-SrtA-$His_{10}$ |
| 11 | *a*MSP9-$L_{14D}$-eSrtA | MSP9-LPGTG-$L_{14D}$-eSrtA-$His_{10}$ |
| 12 | *a*MSP11-$L_{12}$ | MSP11-LPGTG-$L_{12}$-SrtA-$His_{10}$ |
| 13 | *a*MSP11-$L_{14D}$ | MSP11-LPGTG-$L_{14D}$-SrtA-$His_{10}$ |
| 14 | *a*MSP15-$L_{12}$ | MSP15-LPGTG-$L_{12}$-SrtA-$His_{10}$ |
| 15 | *a*MSP15-$L_{14D}$ | MSP15-LPGTG-$L_{14D}$-SrtA-$His_{10}$ |
| 16 | *a*SFTI-$L_{12}$ | SFTI-LPGTG-$L_{12}$-SrtA-His10 |
| 17 | *a*-*i*-SFTI-$L_{19D}$ | Inhibitory Sequence-SFTI-LPGT-$L_{19D}$-SrtA-$His_{10}$ |
| 18 | *a*kB1-$L_{12}$ | kB1-LPVTG-$L_{12}$-SrtA-$His_{10}$ |
| 19 | *a*-*i*-kB1-$L_{19D}$ | Inhibitory Sequence-kB1-LPVT-$L_{19D}$-SrtA-$His_{10}$ |
| 20 | *a*Vc1.1-$L_{12}$ | Vc1.1-LPGTG-$L_{12}$-SrtA-$His_{10}$ |
| 21 | *a*Vc1.1-$L_{19D}$ | Vc1.1-LPGTG-$L_{19D}$-SrtA-$His_{10}$ |
